# A Sodium-dependent Trehalose Transporter Contributes to Anhydrobiosis in Insect Cell Line, Pv11

**DOI:** 10.1101/2023.09.29.560116

**Authors:** Kosuke Mizutani, Yuki Yoshida, Eita Nakanishi, Yugo Miyata, Shoko Tokumoto, Hiroto Fuse, Oleg Gusev, Shingo Kikuta, Takahiro Kikawada

## Abstract

Pv11 is the only animal cell culture that, when preconditioned with a high concentration of trehalose, can be preserved in the dry state at room temperature for more than one year while retaining the ability to resume proliferation. This extreme desiccation tolerance is referred to as anhydrobiosis. Here we identified a novel transporter that contributes to the recovery of Pv11 cells from anhydrobiosis. In general, the SLC5 family of secondary active transporters co-transport Na^+^ and carbohydrates including glucose. Here we identified a novel transporter STRT1 (sodium-ion trehalose transporter 1) belonging to the SLC5 family that is highly expressed in Pv11 cells and transports trehalose with Na^+^ dependency. This is the first report of an SLC5 family member that transports a naturally occurring disaccharide, such as trehalose. Knockout of the *Strt1* gene significantly reduced the viability of Pv11 cells upon rehydration after desiccation. During rehydration, when intracellular trehalose is no longer needed, *Strt1*-knockout cells released the disaccharide more slowly than the parent cell line. During rehydration, Pv11 cells became roughly spherical due to osmotic pressure changes, but then returned to their original spindle shape after about 30 min. *Strt1*-knockout cells, however, required about 50 min to adopt their normal morphology. STRT1 probably regulates intracellular osmolality by releasing unwanted intracellular trehalose with Na^+^, thereby facilitating the recovery of normal cell morphology during rehydration. STRT1 likely improves the viability of dried Pv11 cells by rapidly alleviating the significant physical stresses that arise during rehydration.

**Significance Statement:** This is the first report of an SLC5 family member, STRT1 (sodium ion trehalose transporter 1), with Na^+^-dependent trehalose transport activity. A *Strt1*-knockout cell line revealed that STRT1 likely plays an important role during anhydrobiosis in Pv11 cells: it efficiently discharges unwanted trehalose in the presence of Na^+^ during rehydration of dried Pv11 cells, effectively reducing intracellular osmolality and thereby restoring cell morphology to a normal state.

## Introduction

Some organisms can withstand long periods of extreme desiccation by entering an ametabolic glassy state (1–4). This state, called “anhydrobiosis”, allows organisms to remain apparently lifeless for periods from several months up to two decades (5). When abiotic conditions become more favorable, rehydration enables anhydrobiotic species to resume metabolism and continue their life cycles (6, 7). The largest anhydrobiotic animal known to date is the larva of the sleeping chironomid, *Polypedilum vanderplanki* (7, 8). Pv11, a cell line derived from the egg mass of *P. vanderplanki,* is also capable of anhydrobiosis (9). The genome sequence and transcriptome of *P. vanderplanki* larvae and Pv11 cells has been deciphered (10–13), and several candidate genes have been predicted to be involved in anhydrobiosis. In contrast to *P. vanderplanki* itself, molecular genetic techniques have already been developed for Pv11 cells, including gene transfer and overexpression, as well as gene knockdown and knockout systems (14–17). These allow us to investigate the molecular mechanisms underlying anhydrobiosis in Pv11 cells.

Trehalose, a non-reducing disaccharide, [α-D-glucopyranosyl-(1,1)-α-D-glucopyranoside, Glc(α1–1α)Glc] is one of the key molecules that allows many anhydrobiotic organisms, such as *P. vanderplanki*, nematodes and yeast, to tolerate an ametabolic, dry state while maintaining the ability to return to normal life activities on rehydration (18–20). As the water in the cells of an anhydrobiotic animal evaporates, trehalose fills in to replace the space previously occupied by water in the cells (21). Upon complete desiccation, trehalose forms an organic glass, which is assumed to physicochemically protect all the biological components in the organism (22). Thus, *P. vanderplanki* larvae massively accumulate trehalose as they lose water, which acts as a protectant against desiccation stress (18). To successfully prepare Pv11 cells for anhydrobiosis, a preconditioning step involving incubation in a 600-mM trehalose solution for 48 h is essential prior to desiccation; this step induces anhydrobiosis (23). Following preconditioning and desiccation, Pv11 cells resume proliferation when they are rehydrated by addition of an insect growth medium (23). The preconditioning step that induces anhydrobiosis is likely to involve intracellular accumulation of sufficient trehalose to protect biological components by vitrification as water is lost. On the other hand, during rehydration it is probably essential to release trehalose from the cell as quickly as possible to counteract the influx of water. The presence of excess solutes in the cell leads to an unwanted increase in intracellular osmolality and, in the worst case, to cell lysis and death. Therefore, understanding the cellular trehalose transport system is important in unveiling the molecular mechanisms underlying anhydrobiosis.

Trehalose cannot freely permeate cell membranes, but <u>tre</u>halose transporter <u>1</u> (TRET1, g6534) allows to it successfully pass through (24). TRET1 is a facilitated diffusion-type trehalose transporter belonging to the solute carrier 2 (SLC2) family; such a transporter is common in insects, where it maintains the level of trehalose as the major hemolymph sugar (24, 25). In *P. vanderplanki* and Pv11 cells, trehalose transport via TRET1 is driven by a concentration gradient. TRET1 is thought to play two important roles in anhydrobiosis: it regulates intra- and extracellular osmolarity during dehydration and rehydration; and it delivers trehalose, an essential bioprotectant, into and around cells.

In addition to TRET1, other trehalose transporters have been predicted in Pv11 cells and *P. vanderplanki* larvae. In larvae, an increase in Na^+^ caused by dehydration triggers anhydrobiosis-specific trehalose accumulation, implying that Na^+^ and trehalose are closely related in a physiologically relevant manner in *P. vanderplanki* (26). Previously, desiccation-inducible sodium/solute symporter family (SSF) genes were hypothesized to aid in trehalose transport in peripheral tissues (except the fat body) in desiccating larvae (10). These transporters, which belong to the solute carrier 5 (SLC5) family, are classified as secondary active transporters that transport solutes such as sugars and other carbohydrates and are driven by electrochemical gradients of Na^+^ across cell membranes (27, 28).

In general, transport of sugars such as glucose and trehalose is mediated by members of the SLC superfamily, which comprises more than 50 families, and transports a wide range of solutes (sugars, amino acids, peptides, fats, ions and vitamins) across membranes (29). The SLC2, SLC5, SLC45, SLC50 and SLC60 families have been predicted to be involved in sugar transport (30–36). Two well-characterized SLC families of sugar transporters are SLC2 and SLC5. The SLC2 family genes (Glut1-14, in humans) are reported to facilitate the diffusion of sugars such as glucose across the plasma membrane (37), whereas the SLC5 family genes (SGLT1-6) transport sugars (glucose, fructose, lactate or pyruvate) in a sodium gradient-dependent manner (27, 28, 32). The human SLC5 family consists of 12 genes, all of which are sodium cotransporters, with the exception of SGLT3, which is a glucose sensor (28, 38, 39). Transporters have broad ligand specificity and a single transporter transports multiple substrates (27); for example, SGLT1 mainly cotransports Na^+^ and glucose, but can also cotransport Na^+^ together with urea and water (40, 41). Therefore, it is not possible to predict the substrate to be transported based on the sequence of the protein. In mammals, tissue-specific and temporal regulation of the expression of the SLC2 and SLC5 families underpins an efficient transport system for the blood sugar, glucose (42, 43). As a result, a precise homeostasis of blood glucose is maintained. Insects are thought to have functionally analogous SLC2- and SLC5-mediated systems for sugar metabolism.

To the best of our knowledge, there are no reports that the SLC5 family is involved in the transport of trehalose. To date in insects, besides TRET1, three other proteins, HaST10, Pippin and MFS3, have been reported to transport trehalose (24, 25, 44-47). TRET1, HaST10 and Pippin are all facilitative diffusion-type passive transporters belonging to the SLC2 family (24, 25, 44-47). MFS3 belonging to the SLC17 family transports trehalose and glucose (46). In the fruit fly, *Drosophila melanogaster*, 14 SLC5 proteins are encoded in the genome (48). Only three of these genes have been investigated in detail but, while *Drosophila* SLC5 family members can transport glucose (49–52), it is not yet known whether they transport trehalose.

In this study, we found that the protein encoded by *g4064*, which we named <u>s</u>odium-ion <u>tr</u>ehalose transporter <u>1</u> (STRT1), is a member of the SLC5 family in *P. vanderplanki* and has trehalose transport activity. Disruption of this gene indicates that STRT1 plays an important role in anhydrobiosis in Pv11 cells.

## Results

### Knockout of *g4064* impairs anhydrobiosis in Pv11 cells

We searched for genes encoding proteins with SLC5 (SSF)-specific motifs in the latest version of the *P. vanderplanki* genome (10) and identified 22 examples (Table 1). We then compared the expression profiles of five of the 22 genes that were significantly expressed in Pv11 cells (Fig. 1A and B, Supplemental Dataset S1). Of these, *g4064* was the most highly expressed and its expression was upregulated during rehydration after anhydrobiosis as the cells returned to normal physiological conditions.

**Figure 1.**
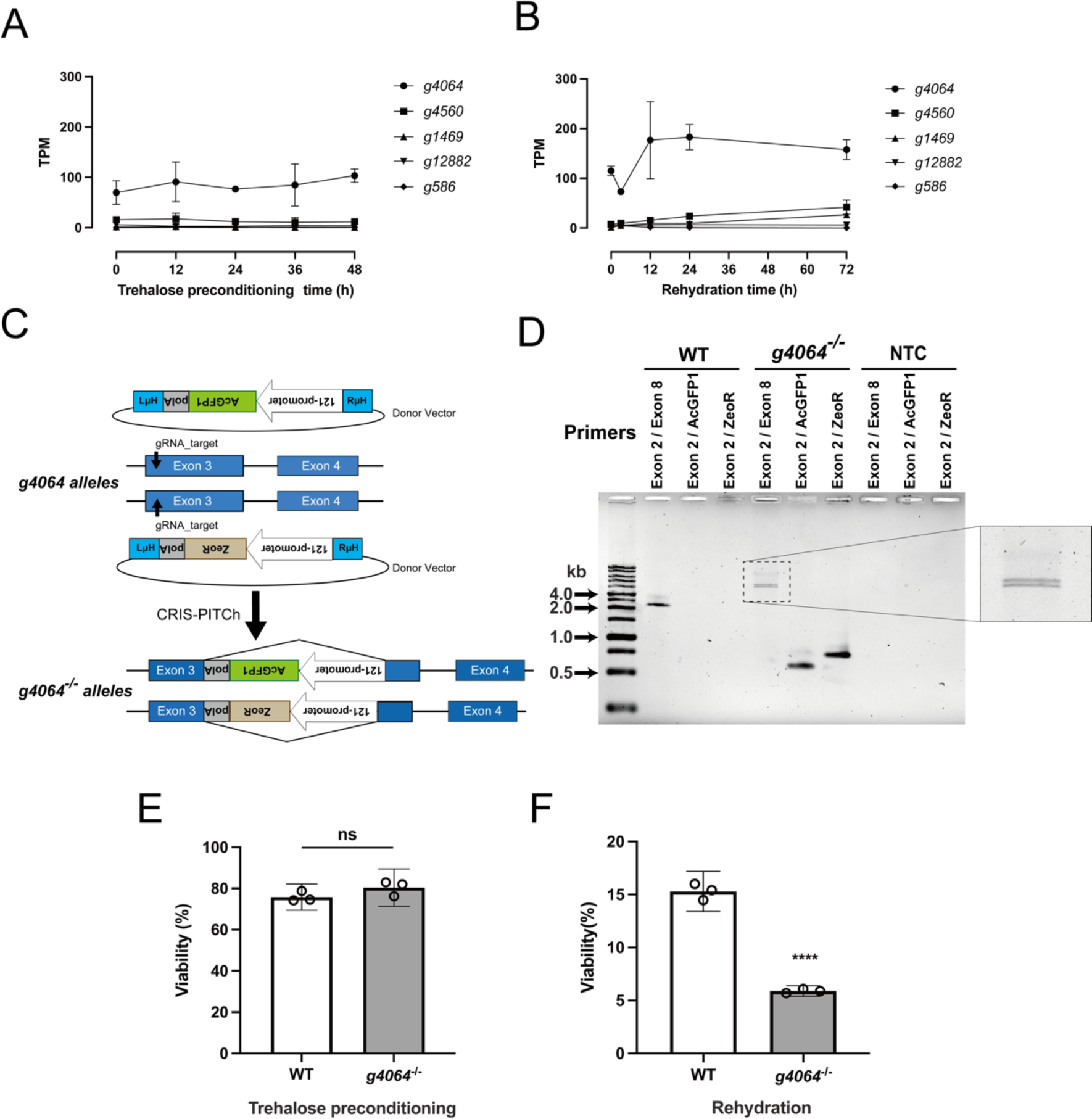
Disruption of an SLC5 gene, *g4064*, which is highly expressed after rehydration, impacts anhydrobiosis in Pv11 cells. From the SLC5 family genes of *P. vanderplanki*, we selected those with an average mRNA expression level >1 TPM during preconditioning (**A**) and rehydration (**B**) in Pv11 cells. For other expression data, see Supplemental Dataset S1. (**C**) Schematic diagram for the establishment of a *g4064*-knockout (*g4064*^−/−^) cell line from Pv11 cells using the CRIS-PITCh system. Donor vectors harboring the expression units for either AcGFP1 or ZeoR flanked by microhomology arms (μH) were co-transfected with gRNA- and hSpCas9-expression vectors into Pv11 cells, resulting in the insertion of either the AcGFP1 or the ZeoR expression unit into exon 3 of each allele of *g4064*. (**D**) Genomic PCR was conducted to confirm the insertion of exogenous expression units into the *g4064* alleles in the knockout cells (*g4064*^−/−^) in comparison to the intact alleles in Pv11 cells (WT). (**E**) Viability after trehalose preconditioning of WT cells and the *g4064*^−/−^ clonal cell line. Cell viability was determined after 48 h of trehalose preconditioning. “ns” indicates no significant difference. (**F**) Viability after desiccation and 24 h rehydration of WT cells and the *g4064*^−/−^ clonal cell line. Tukey’s test result shows ****: *p* ≤ 0.0001. Data indicate mean ± 95% CI, n = 3 in each group.

**Table 1.** SLC5 family genes of *P. vanderplanki* identified by hmmsearch.

| Gene ID | Representative protein | Domain | Start | End | Accession_Pfam | Score | E-value | Description |
| --- | --- | --- | --- | --- | --- | --- | --- | --- |
| g4064 | KAG5674078.1 | SSF | 38 | 449 | PF00474.20 | 132.5 | 2.00E-38 | Sodium:solute symporter family |
| g4560 | KAG5674608.1 | SSF | 167 | 452 | PF00474.20 | 63.6 | 1.60E-17 | Sodium:solute symporter family |
| g1469 | KAG5671264.1 | SSF | 43 | 450 | PF00474.20 | 86.8 | 1.40E-24 | Sodium:solute symporter family |
| g12882 | KAG5683611.1 | SSF | 104 | 508 | PF00474.20 | 100.8 | 8.10E-29 | Sodium:solute symporter family |
| g586 | KAG5670311.1 | SSF | 55 | 460 | PF00474.20 | 137.3 | 6.80E-40 | Sodium:solute symporter family |
| g6015 | KAG5676168.1 | SSF | 51 | 458 | PF00474.20 | 141.7 | 3.10E-41 | Sodium:solute symporter family |
| g12607 | KAG5683323.1 | SSF | 38 | 449 | PF00474.20 | 134 | 6.60E-39 | Sodium:solute symporter family |
| g5003 | KAG5675068.1 | SSF | 51 | 455 | PF00474.20 | 127 | 8.90E-37 | Sodium:solute symporter family |
| g12037 | KAG5682705.1 | SSF | 68 | 472 | PF00474.20 | 129.6 | 1.50E-37 | Sodium:solute symporter family |
| g1791 | KAG5671599.1 | SSF | 38 | 431 | PF00474.20 | 108.5 | 3.90E-31 | Sodium:solute symporter family |
| g16375 | KAG5668440.1 | SSF | 78 | 222 | PF00474.20 | 55.6 | 4.40E-15 | Sodium:solute symporter family |
| g7082 | KAG5677315.1 | SSF | 38 | 449 | PF00474.20 | 129.3 | 1.80E-37 | Sodium:solute symporter family |
| g8951 | KAG5679385.1 | SSF | 122 | 526 | PF00474.20 | 108.5 | 3.80E-31 | Sodium:solute symporter family |
| g3876 | KAG5673866.1 | SSF | 48 | 454 | PF00474.20 | 141.2 | 4.50E-41 | Sodium:solute symporter family |
| g14930 | KAG5666928.1 | SSF | 61 | 463 | PF00474.20 | 111.9 | 3.40E-32 | Sodium:solute symporter family |
| g3434 | KAG5673379.1 | SSF | 55 | 459 | PF00474.20 | 134.6 | 4.50E-39 | Sodium:solute symporter family |
| g16377 | KAG5668442.1 | SSF | 91 | 420 | PF00474.20 | 74.6 | 7.40E-21 | Sodium:solute symporter family |
| g10696 | KAG5681247.1 | SSF | 69 | 476 | PF00474.20 | 147.6 | 5.10E-43 | Sodium:solute symporter family |
| g10695 | KAG5681246.1 | SSF | 76 | 483 | PF00474.20 | 140.1 | 9.90E-41 | Sodium:solute symporter family |
| g16374 | KAG5668439.1 | SSF | 63 | 465 | PF00474.20 | 107.8 | 6.30E-31 | Sodium:solute symporter family |
| g16376 | KAG5668441.1 | SSF | 63 | 137 | PF00474.20 | 26.4 | 3.20E-06 | Sodium:solute symporter family |
| g8949 | KAG5679383.1 | SSF | 6 | 142 | PF00474.20 | 24.3 | 1.40E-05 | Sodium:solute symporter family |

To investigate the physiological function of *g4064* (cDNA and genome structure: Supplemental Dataset S2) in Pv11 cells, exon 3 of each allele of *g4064* was disrupted by insertion of expression units for AcGFP1 and ZeoR sequences, respectively, using CRIS-PITCh (CRISPR/Cas9-mediated precise integration into target chromosome) (Fig. 1C, Supplemental Fig. S1) (16, 53). The cell line carrying the exogenous genes exhibited strong AcGFP1 fluorescence (Supplemental Fig. S2). Genomic DNA was extracted from these cells and PCR with specific primers for the *g4064* genes confirmed both insertions of exogenous DNA (Fig. 1D). The sequence of the products amplified from the CRIS-PITCh-treated cells indicated that the AcGFP1 and ZeoR expression units were inserted into exon 3 of each allele of *g4064* as expected (Supplemental Dataset S2). These results demonstrated the successful establishment of a biallelic *g4064*-knockout Pv11 cell line (*g4064*^−/−^). Under normal conditions, there was no significant difference in proliferation rates between wild-type Pv11 (WT) and *g4064*^−/−^ cells (Supplemental Fig. S3). Similarly, 48 h after trehalose pretreatment, there was no significant difference in the viability of WT and *g4064*^−/−^ cells (Fig. 1E). However, the viability of *g4064*^−/−^ cells after desiccation and rehydration was significantly lower than that of WT cells (Fig. 1F), suggesting that *g4064* contributes to the process of anhydrobiosis in Pv11 cells.

### *g4064* encodes a protein with trehalose transport activity

To investigate whether the g4064 protein has the ability to transport trehalose, we expressed *g4064* heterologously in CHO-K1 cells and *Xenopus* oocytes. First, we measured the transport activity of the g4064 protein in CHO-K1 cells. We generated CHO-*g4064*-expressing cells and empty-vector transfected cells (CHO-X cells, a negative control) using the Flip-In system (Thermo Fisher Scientific). Note that CHO-X cells were able to import a non-metabolizable glucose analog, 2-deoxy-glucose (2-DOG), but not trehalose, into their cytoplasm, regardless of salinity conditions (Fig. 2A and B). On the other hand, both glucose and trehalose uptake were elevated in CHO-*g4064*-expressing cells under high-salinity conditions (Fig. 2A and B). To examine the substrate selectivity of the g4064 protein as a sugar transporter, we conducted uptake assays for trehalose, maltose, sucrose, lactose and cellobiose using the *Xenopus* oocyte expression system. We found that *g4064*-expressing oocytes incorporated trehalose to a significant degree (Fig. 2C). The kinetics of trehalose transporter activity of *g4064*-expressing oocytes were measured under high-salinity conditions (100 mM NaCl), yielding a Km value of 265.4 mM and a Vmax of 69.1 pmol/min/oocyte (Fig. 2D).

**Figure 2.**
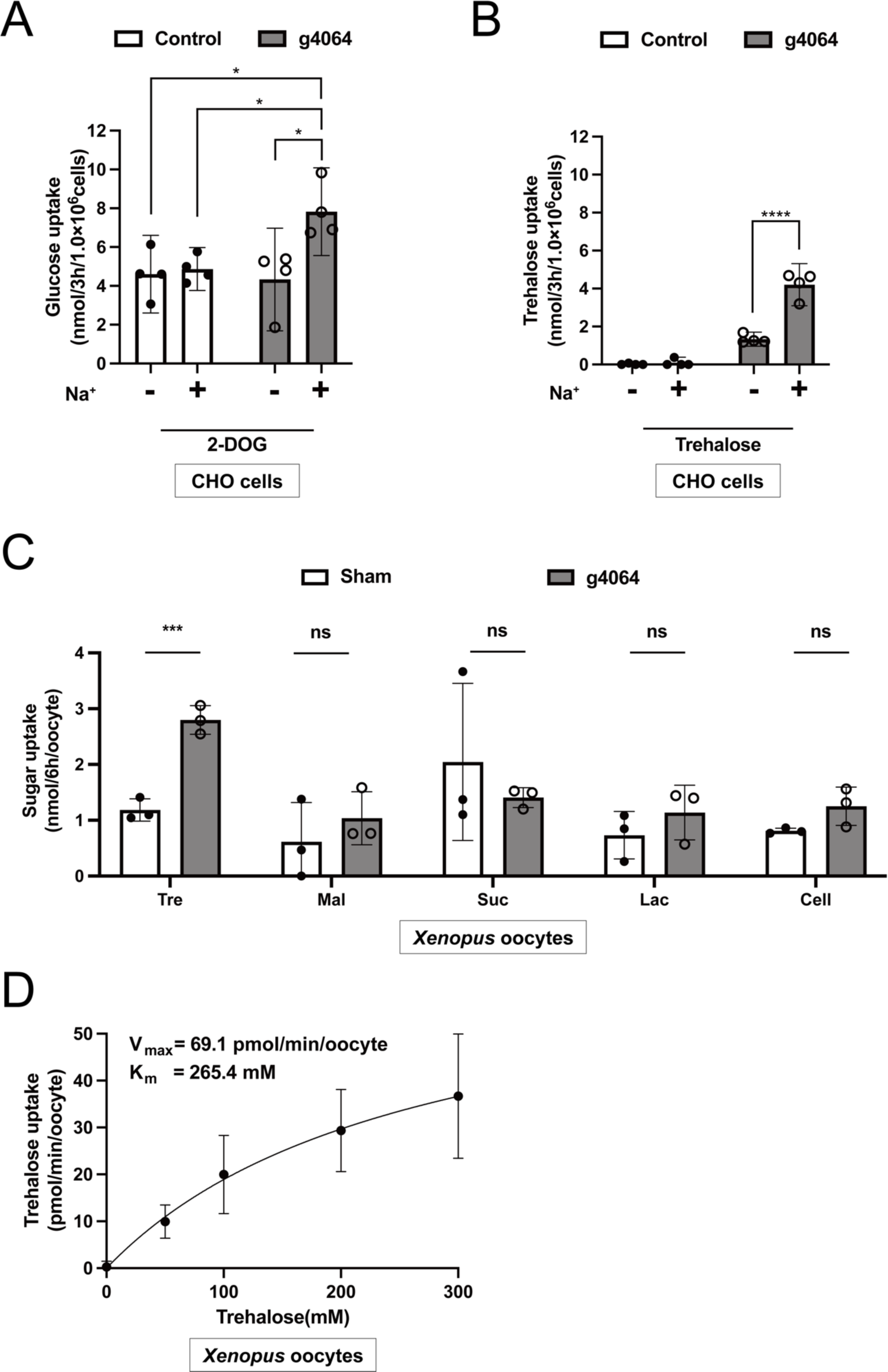
The *g4064* gene encodes a trehalose transporter. **(A)** Glucose uptake in CHO cells expressing *g4064*. The Na^+^-containing buffer solution (+) comprised 100 mM 2-DOG and 100 mM NaCl in HBSS buffer, while the Na^+^-limiting buffer solution (–) was identical except that NaCl was replaced with choline chloride. Control: CHO-X cells; g4064: CHO-*g4064*-expressing cells. Tukey’s test result shows *: *p* ≤ 0.05. (**B**) Trehalose uptake via CHO cells expressing *g4064*. The Na^+^-containing buffer solution (+) comprised 100 mM trehalose and 100 mM NaCl in HBSS buffer, while the Na^+^-limiting buffer solution (–) was identical except that NaCl was replaced with choline chloride. Tukey’s test result shows ****: *p* ≤ 0.0001. (**C**) Disaccharide selectivity of the g4064 protein expressed in *Xenopus* oocytes. Tre, trehalose; Mal, maltose; Suc, sucrose; Lac, lactose; Cel, cellobiose. Tukey’s test result shows ***: *p* ≤ 0.001. (**D**) Kinetics of the g4064 protein expressed in *Xenopus* oocytes for trehalose uptake. The uptake assays were performed using a buffer containing 200 mEq/L Na^+^. Data indicate mean ± 95% CI, n = 3-4 in each group.

### The g4064 protein is a Na^+^-dependent trehalose transporter

Since the g4064 protein belongs to the SLC5 family, we assumed that this trehalose transporter should transport trehalose along a Na^+^ electrochemical gradient. Accordingly, trehalose influx due to the g4064 protein increased in a Na^+^-dependent manner (Fig. 3A). We analyzed trehalose uptake in *g4064*-expressing oocytes under Na^+^-limiting conditions and found a decrease in trehalose influx when NaCl in the *Xenopus* expression system was replaced by an equivalent concentration of choline chloride (Fig. 3B). We next examined the possibility of counter anion (Cl^-^)-dependent transport, but found that *g4064*-expressing oocytes in Cl^-^-free buffers containing either Na_2_SO_4_ or NaNO_3_ showed essentially identical trehalose uptake activity as oocytes in Cl^-^-containing buffers (Fig. 3B). Phlorizin is known to act as a competitive inhibitor of SLC5 family members, such as SGLT1 and SGLT2 (54). The trehalose-transport activity of the g4064 protein was significantly inhibited by phlorizin, confirming that its biochemical activity is consistent with it being a member of the SLC5 family (Fig. 3C). In general, the activity of secondary active transport membrane transporters, like the SLC5 family, is driven by electrochemical membrane gradients of ions, such as Na^+^ and H^+^. To investigate whether g4064 protein activity is driven by H^+^, its trehalose transport activity was examined at various pH levels, but no significant change was recorded (Fig. 3D). These results revealed that the g4064 protein is a Na^+^-dependent trehalose transporter, and consequently we named it sodium-ion trehalose transporter 1 (STRT1).

**Figure 3.**
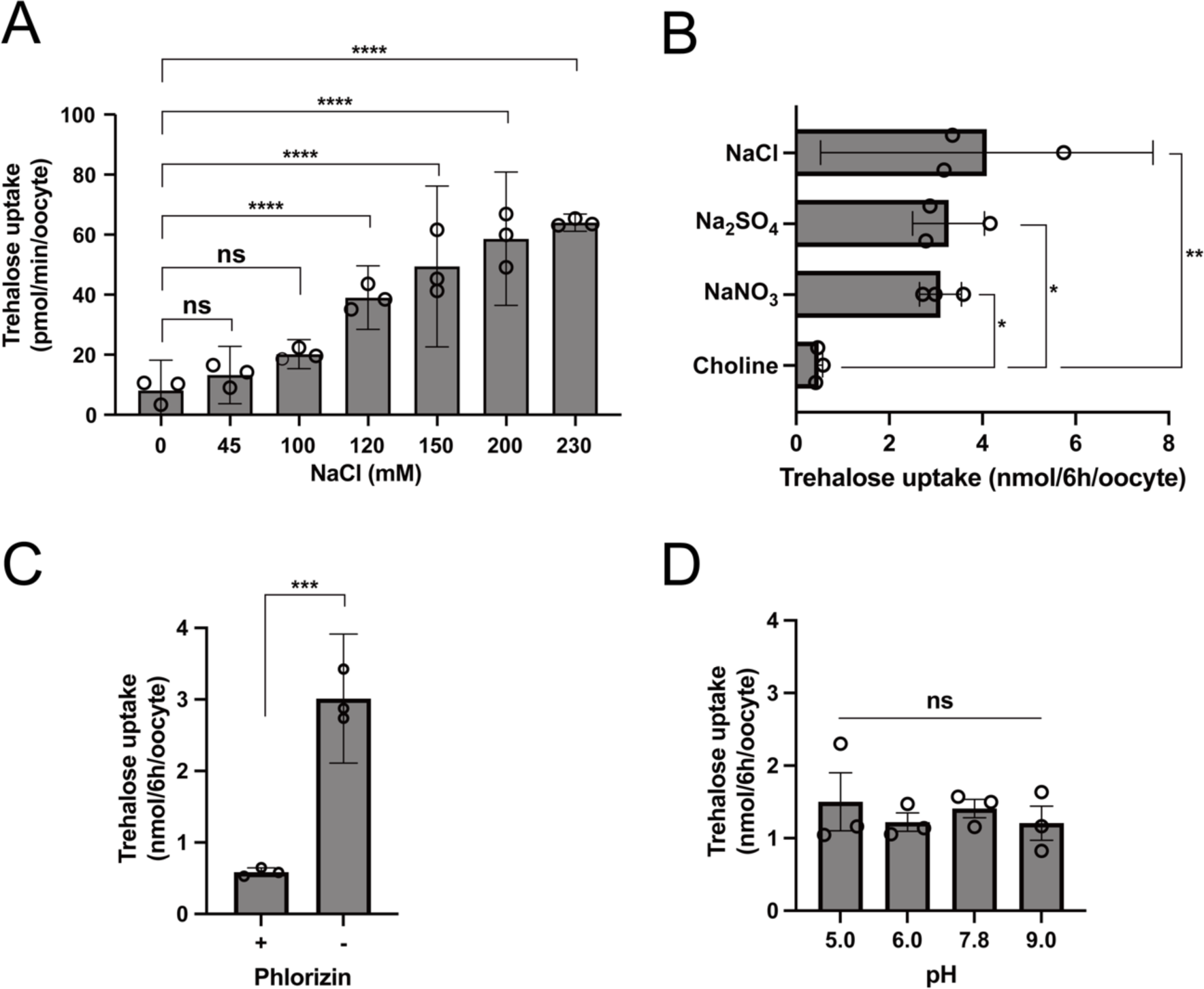
Extracellular Na^+^ enhances trehalose transport activity of the g4064 protein. Trehalose uptake assays were examined in *g4064*-expressing *Xenopus* oocytes. Twenty oocytes were analyzed in each assay. **(A)** Dose response of trehalose uptake by the g4064 protein at various Na^+^ ion concentrations. Dunnett’s test result shows as “ns” indicates no significant difference (*p* > 0.05); ****: *p* ≤ 0.0001. **(B)** Na^+^ dependency of trehalose uptake by STRT1. Choline chloride (‘choline’) indicates a Na^+^-limiting buffer solution was used. Tukey’s test result shows as *: *p* ≤ 0.05, **: *p* ≤ 0.01. **(C)** Inhibition by 100 µM phlorizin. The uptake assays were performed using a buffer containing 200 mEq/L Na^+^. Tukey’s test result shows ***: *p* ≤ 0.001. **(D)** pH dependency of trehalose uptake by the g4064 protein. Ten oocytes were analyzed in each assay. Trehalose transport activity was examined in MBS buffer containing 2-(N-morpholino) ethanesulfonic acid (MES) at pH 5.0–6.0 or in Tris HCl at pH 7.8–9.0. Tukey’s test result shows “ns” indicates no significant difference (*p* > 0.05). ****: *p* ≤ 0.0001. Data indicate mean ± 95% CI, n = 3 in each group.

### STRT1 contributes to the efficient efflux of trehalose during rehydration

To confirm that the trehalose transport activity of STRT1 is Na^+^-dependent in Pv11 cells, we examined trehalose uptake in the presence of various NaCl concentrations. A significant increase in trehalose influx with increasing NaCl concentration was observed in WT but not in a *Strt1-*knockout cell line (*Strt1*^−/−^ = *g4064*^−/−^) (Fig. 4A). Thus, STRT1 indeed functions in Pv11 cells as a Na^+^-dependent trehalose transporter.

**Figure 4.**
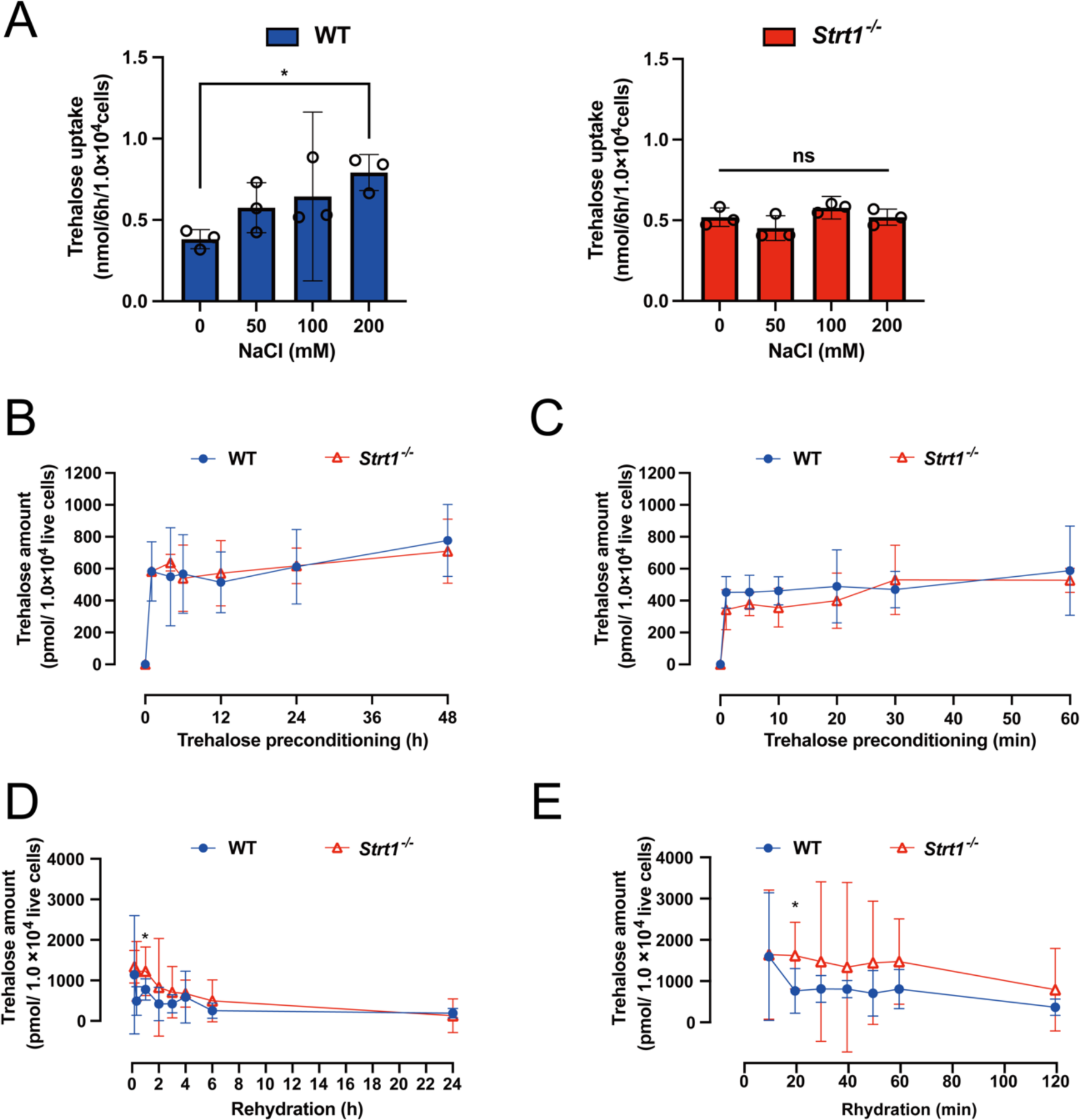
Comparison of trehalose content in *Strt1^−/−^* and WT cells. **(A)** Disruption of *Strt1* causes loss of NaCl-dependent trehalose transport activity in Pv11 cells. Uptake assays were performed in buffers containing various concentrations of NaCl with 400 mM trehalose. Tukey’s test result shows *: *p* ≤ 0.05; “ns” indicates no significant difference (*p* > 0.05). **(B)** Changes in trehalose content in cells during preconditioning with 600 mM trehalose solution as the induction step for anhydrobiosis. **(C)** Focusing on the 60-min period immediately following preconditioning to obtain information on changes in trehalose content at more precise time intervals. **(D)** Changes in cellular trehalose content after desiccation and rehydration. Tukey’s test result shows *: *p* ≤ 0.05. **(E)** Focusing on the 60-min period immediately following rehydration, to obtain information on changes in trehalose content at more precise time intervals. Tukey’s test result shows *: *p* ≤ 0.05. Data indicate mean ± 95% CI, n = 3-4 in each group.

We next examined the involvement of STRT1 in the regulation of intracellular trehalose concentration during the trehalose pretreatment and rehydration steps in Pv11 cells. There was no significant difference between WT and *Strt1*^−/−^ in trehalose uptake capacity during preconditioning (Fig. 4B). Trehalose uptake in Pv11 cells was at its maximum at about 6 h, so we examined the early time points. Even at the 1 h time point, when rapid trehalose uptake appears to be taking place, there was no significant difference between WT and *Strt1*^−/−^ cells in trehalose uptake capacity (Fig. 4C). Upon rehydration, intracellular trehalose was rapidly released from WT cells, but this trehalose efflux was delayed in the *Strt1*^−/−^ knockout line. As a result, the amount of intracellular trehalose had decreased significantly faster in WT than in *Strt1*^−/−^ cells after 20 min rehydration. Thereafter (30 min after rehydration), however, no significant differences were observed between the two cell types (Fig. 4D). Analysis of the efflux of trehalose during the early stages of rehydration also only showed a significant difference after 20 min of rehydration (Fig. 4E). These data show that STRT1 contributes to trehalose efflux in Pv11 cells after rehydration.

### STRT1 promotes the recovery of normal morphology in Pv11 cells on rehydration

Pretreatment with a highly concentrated trehalose solution, followed by desiccation and rehydration, leads to enormous changes in osmotic pressure in Pv11 cells, which will impact cell morphology. To analyze the changes in the morphology of live cells during anhydrobiosis, AcGFP1-positive Pv11 cell lines were analyzed. We used *Strt1*^−/−^ cells and, as a control, an *AcGFP1*^+^*/ ZeoR*^+^ cell line in which the exogenous genes AcGFP1 and ZeoR were inserted into a safe harbor region of the genome (chromosome 1: intergenic region between g1212124 and g1212125) (55). The *AcGFP1*^+^*/ ZeoR*^+^ cells showed comparable desiccation tolerance to WT (55).

Morphological changes in *Strt1*^−/−^ and *AcGFP1*^+^*/ ZeoR*^+^ cells during trehalose preconditioning and rehydration were examined. Both cell lines showed no difference in cell circularity under normal culture conditions in IPL-41 medium, when Pv11 cells are spindle-shaped. Pretreatment with trehalose increased the cell circularity of both cell lines, but there was no significant difference between them (Fig. 5A, Fig. S4). Within 10 min after rehydration was initiated, the circularity of both cell types increased slightly due to swelling caused by the difference in osmolality between the inside and outside of the cells (Fig. 5B, Fig. S5). Thereafter, by 40 min after rehydration, the circularity of *AcGFP1*^+^*/ ZeoR*^+^ cells returned to normal levels, whereas the cell circularity of the *Strt1*^−/−^ line took about 60 min to normalize (Fig. 5B, Fig. S5). The viability of both cell lines was not significantly different at 10 min after rehydration. However, the viability of *Strt1*^−/−^ decreased significantly compared to *AcGFP1*^+^*/ ZeoR*^+^ cells from 30 min onwards (Fig. 5C).

**Figure 5.**
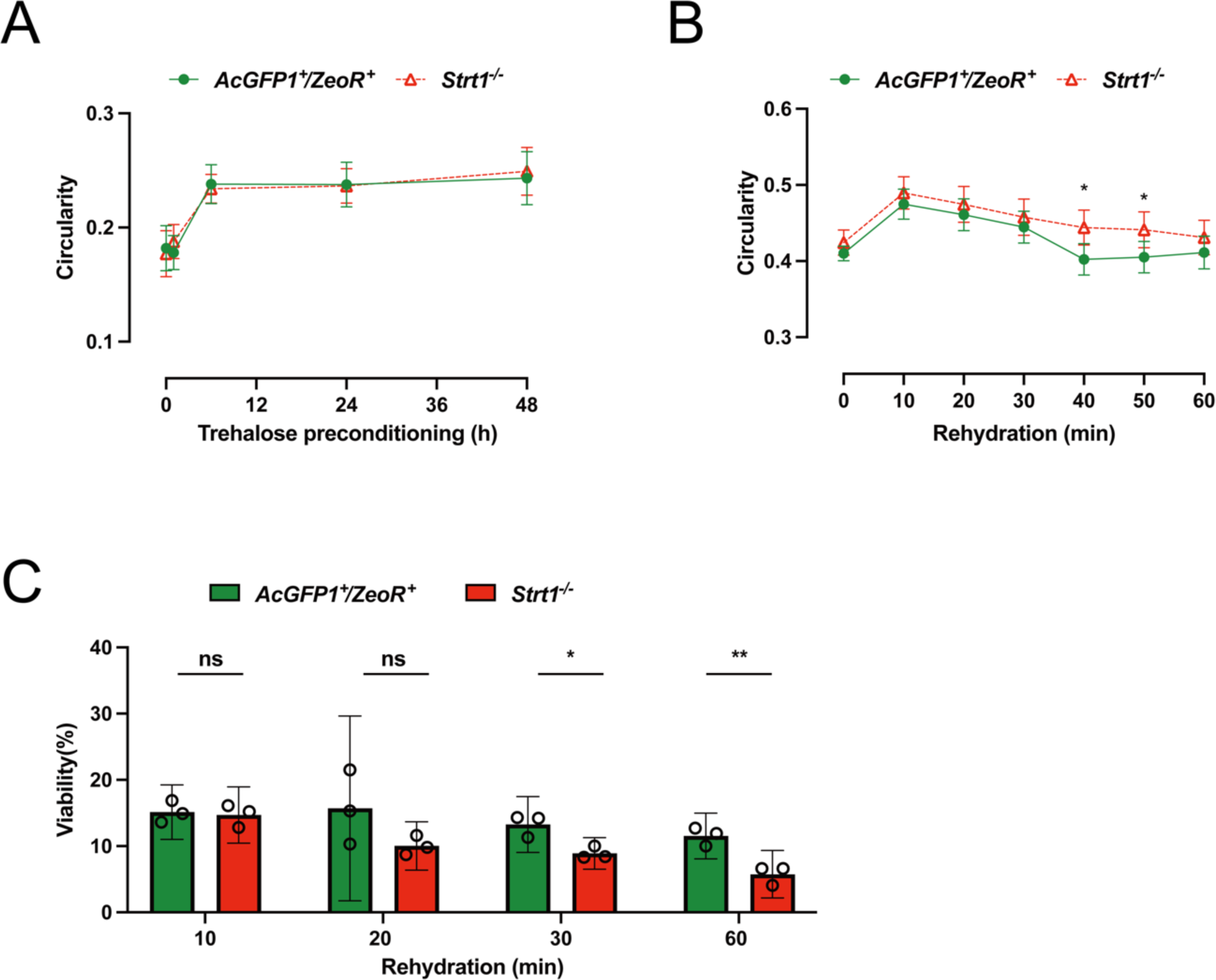
Changes in cell circularity of Pv11 cells during anhydrobiosis. Comparison of changes in cell circularity between *AcGFP1*^+^/*ZeoR* ^+^ and *Strt1*^−/−^cells. Over 100 cells were used for the analysis. Changes in circularity of the cells **(A)** during preconditioning and **(B)** after rehydration; Tukey’s test result shows *: *p* ≤ 0.05. Data indicate mean ± 95 % CI, n ≥ 100 in each group. **(C)** Viability of cells after desiccation and rehydration. Tukey’s test result shows “ns” indicates no significant difference (*p* > 0.05), *: *p* ≤ 0.05, **: *p* ≤ 0.01. Data indicate mean ± 95% CI, n = 3 in each group.

## Discussion

In this study, we demonstrated that STRT1 exhibits Na^+^-dependent trehalose transport activity in Pv11 cells. Moreover, *Strt1*^−/−^ cells exhibited lower viability after desiccation and rehydration than intact Pv11 cells (Fig. 1E, F). *Strt1*^−/−^ cells were significantly slower to discharge redundant intracellular trehalose upon rehydration than WT, causing a delay in the return to normal morphology after rapid osmotic swelling (Fig. 5A, B). STRT1 is likely to be responsible for the efficient efflux of unwanted trehalose during rehydration in Pv11 cells, which is required to effectively decrease intracellular osmotic pressure and thereby quickly restore cell morphology to its normal state.

Heterologous expression systems showed that trehalose transport by STRT1 requires a Na^+^ gradient (Fig. 2 and 3). In *Xenopus* oocytes overexpressing STRT1, Na^+^-dependent trehalose uptake was inhibited by phlorizin, an inhibitor of SGLT1 and SGLT2 (Fig. 2 and 3) (54). STRT1 can be classified as a secondary active transporter of the SLC5 family. In general, secondary active transporters allow the transport of substrates across the cellular membrane against the substrate concentration; this is driven by the electrochemical concentration gradient of Na^+^ (27, 28, 32). The human SLC5 family genes encode sodium cotransporters, except for SGLT3 (28, 38, 39); for example, SGLT1 and SGLT2 cotransport Na^+^ and glucose (40, 41). Therefore, STRT1 may cotransport Na^+^ and trehalose, like SGLT1 and SGLT2, or transport trehalose using a Na^+^ concentration gradient in some other way; the precise transport mechanism needs to be experimentally confirmed.

STRT1 possesses trehalose transport activity, but does not seem to be the main route of trehalose import during preconditioning (Fig. 4B, C): the preconditioning buffer contains about 5 mM Na^+^, which is probably too low for STRT1 to perform its function optimally. Strikingly, immediately after rehydration, intracellular trehalose concentrations decreased faster in WT than in *Strt1*^−/−^ cells (Fig. 4D, E). Due to the water loss associated with desiccation, intracellular ion concentrations are expected to increase. STRT1 may respond to changes in these ion concentrations as well as transporting trehalose. The SLC5 family normally transports sodium and substrates into the cell using intra- and extracellular concentration gradients, where the extracellular Na^+^ concentration is usually higher than the intracellular Na^+^ concentration (27, 28, 32). In contrast, during rehydration, the intracellular sodium concentration in Pv11 cells may be elevated compared to normal conditions due to the desiccation step previously. Therefore, STRT1 may transport trehalose together with sodium or use a concentration gradient. This clearly indicates that STRT1 is associated with the extracellular release of trehalose during rehydration.

The facilitated diffusion-type trehalose transporter, TRET1, may also be involved in anhydrobiosis in Pv11 cells. Previously, we showed that TRET1 is likely involved in trehalose accumulation during the induction of anhydrobiosis in *P. vanderplanki* (24). Indeed, *Tret1* is expressed at physiologically functional levels in Pv11 cells (Supplemental Dataset S3). Kinetic analysis revealed that *P. vanderplanki* TRET1 exhibits a high substrate capacity for trehalose with a Km of approximately 115 mM and a very fast trehalose transport rate with a Vmax of 520 pmol/min/cell in *Xenopus* oocytes (24). The kinetic properties of STRT1 indicate a substrate affinity (Km) of about 265 mM for trehalose, which is more than twice as high as that of TRET1, and a Vmax of about 70 pmol/min/cell in *Xenopus* oocytes (24), which is only about 1/7.5 of the maximum trehalose transport rate of TRET1 (Fig. 3C). This difference in kinetics as a trehalose transporter might explain why no difference in trehalose uptake occurred during preconditioning even when *Strt1* function was completely disrupted. As mentioned above, the trehalose concentration of 600 mM during preconditioning, which is required for trehalose uptake into Pv11 cells, is extremely high (23), far exceeding the Km values of both TRET1 and STRT1. In general, Km is an indicator of the optimal substrate concentration for enzyme and transporter activities (56). Although the Km of STRT1 is closer to the trehalose concentration used in preconditioning than that of TRET1, it seems likely that trehalose uptake during preconditioning is almost exclusively through TRET1 because of the overwhelmingly higher trehalose transport rate (i.e., Vmax) of TRET1. The fact that a decrease in intracellular trehalose was also observed in *Strt1*^−/−^ cells means that it is still possible that other transporters are involved in this process. Hence, we attempted to understand the role of TRET1 in Pv11-cell anhydrobiosis by generating a *Tret1*-knockout cell line. However, we were unable to obtain *Tret1*^−/−^ cells by CRIS-PITCh (Supplementary text). This implies that TRET1 might not only regulate trehalose transport during anhydrobiosis, but also play an essential role in normal cell proliferation in Pv11 cells.

*Strt1*^−/−^ cells had a reduced capacity to restore normal cell morphology during the rehydration process. Typically, Pv11 cells are spindle-shaped (9). During trehalose preconditioning and rehydration, Pv11 cells lose this morphology and become markedly more spherical (Supplemental Fig. S4 and S5). Cell circularity was therefore used to quantify cell recovery during trehalose preconditioning and rehydration (Fig. 5A and B). Ten minutes after rehydration, both *AcGFP1*^+^*/ ZeoR*^+^ and *Strt1*^−/−^ cells displayed an increase in cell circularity (i.e., closer to 1), suggesting that significant cell stress had occurred (Fig. 5B). Thereafter, *AcGFP1*^+^*/ ZeoR*^+^ cells significantly decreased in circularity (i.e., were returning to a more normal spindle morphology), while the circularity of *Strt1*^−/−^ cells remained significantly higher (Fig. 5B). The rehydration process involves a rapid change in osmotic pressure and the rapid influx of water into the cells is likely to cause significant structural stress to the cellular membrane and intracellular structures. Cells, including those of bacteria and humans, monitor cellular conditions such as osmotic pressure and regulate cell shape and water content in response to osmotic stress (57, 58). The response mechanism differs between cell types, but the transport of water, electrolytes (Na^+^, K^+^ and Cl^-^ ions) and organic osmolytes (sorbitol, inositol, glycerophosphorylcholine, betaine, taurine, trehalose and amino acids) is used to adapt the cytoplasmic osmotic pressure to the surrounding environment (59–63). The efflux activity of STRT1 may alleviate osmotic stress by regulating the changes in intracellular and extracellular concentrations of trehalose and Na^+^ that may occur during desiccation and rehydration. Furthermore, STRT1 may also help to restore a normal Na^+^ electrochemical concentration gradient in rehydrated Pv11 cells. Unfortunately, to the best of our knowledge, there are currently no methods available to quantify intracellular Na^+^ levels in dried and immediately rehydrated cells, and consequently it is difficult to test the above speculations. Considering that dried *Strt1*^−/−^ cells showed significantly reduced viability after rehydration (Fig. 5C), we conclude that STRT1 promotes the recovery of cellular morphology upon rehydration of Pv11 cells by discharging unwanted trehalose, and thereby effectively reviving the dried cells.

STRT1 has the ability to transport glucose (Fig. 2A) and in this role it might also contribute to desiccation tolerance. Since trehalose is a dimer of glucose, transporters that transport trehalose are very likely to be able to transport glucose as well. Indeed, trehalose transporters such as TRET1, Pippin, and MFS3 also exhibit glucose transport activity (24, 25, 44-47). In Pv11 cells, trehalose imported into the cells during preconditioning is partially degraded to glucose just after rehydration by the activity of trehalase (Treh), a trehalose-degrading enzyme. During trehalose preconditioning, there was no significant difference in the amount of intracellular glucose in WT and *Strt1*^−/−^ cells (Fig. S6A). After rehydration, glucose levels in *Strt1*^−/−^ cells became significantly greater than in WT cells at 20 min after rehydration, but then there was no difference in intracellular glucose concentration 1 h after rehydration (Fig. S6B). Any glucose produced should be catalyzed to glucose-6-phosphate (G-6-P) by hexokinase (Hex). G-6-P is known as an initial substrate for glycolysis and the pentose-phosphate pathway (PPP). Naturally, glycolysis contributes to ATP production in the cells. The PPP is activated during rehydration of dried *P. vanderplanki* larvae, leading to synthesis of NADPH, which acts to drive the antioxidant system that alleviates the toxic effects of reactive oxygen species generated during rehydration (64). In Pv11 cells, Treh and Hex were highly expressed during trehalose pretreatment and rehydration (Supplement-Dataset S3) (13). In *P. vanderplanki* larvae, Treh expression is elevated just before desiccation, but it is not enzymically active. Upon rehydration, Treh is quickly activated and starts degrading trehalose (64, 65). This suggests that after rehydration Pv11 cells, like the larvae, may activate the PPP to achieve a successful recovery from anhydrobiosis. The synthesis of two molecules of glucose from one molecule of trehalose can lead to an increase in osmolarity. Thus, although the degradation of trehalose to glucose is essential for the resumption of cellular activity after anhydrobiosis, its intracellular concentration must be regulated so that it does not cause cell lysis. STRT1 could play an important role during rehydration by removing excess trehalose and glucose from the cells. Of course, STRT1 is not the only osmotic regulator in Pv11 cells during the recovery process. In fact, even in *Strt1^−/−^* cells, the rate of decrease in intracellular trehalose concentration was only somewhat retarded and the gene knockout did not cause widespread cell death.

In this study, we demonstrated the importance of STRT1 in Pv11-cell anhydrobiosis. The role of STRT1 in the anhydrobiosis of *P. vanderplanki* larvae also needs to be elucidated. Genomic information shows that there are still uncharacterized transporters in *P. vanderplanki*, some of which may be trehalose transporters. Future identification of additional novel trehalose transporters and elucidation of their functions may lead to a better molecular understanding of anhydrobiosis.

## Materials and Methods

### Identification of SLC5 family genes in *P. vanderplanki* genome

The genome, gene predictions, annotations (accession number GCA_018290095.1) and RNA-Seq data (accession number GSE158443) from our previous study were downloaded from NCBI (13). Gene expression was quantified using RSEM v1.3.1 (--bowtie2) (66) and the following downstream analysis was performed using the Trinity package v2.15.1 (abundance_estimates_to_matrix.pl, run_DE_analysis.pl) (67). Transcripts having FDR < 0.05 were determined as differentially expressed. To identify SSF orthologs, we submitted the amino acid sequences to hmmsearch (--domE 1e-3) (68). The HMM profile for PF00474.20 (sodium/solute symporter family; SSF) was downloaded from InterPro (69).

### Cell culture

Pv11 cells and CHO cells were grown as previously described (9, 18, 70, 71). Pv11 cells were cultured in IPL-41 medium (Thermo Fisher Scientific, Waltham, MA, USA) supplemented with 2.6 g/L tryptose phosphate broth (Becton, Dickinson and Company, Franklin Lakes, NJ, USA), 10% (v/v) fetal bovine serum and 0.05% (v/v) of an antibiotic and antimycotic mixture (penicillin, amphotericin B, and streptomycin; Sigma–Aldrich, St. Louis, MO, USA), designated hereafter as complete IPL-41 medium. A CHO-K1 derivative cell line, Flip-In^TM^-CHO cells (purchased from Thermo Fisher Scientific), was cultured in Ham’s F-12 medium (Sigma–Aldrich) containing 10% FBS (Tissue Culture Biologicals, Tulare, CA), 100 units/ml penicillin and 100 μg/ml streptomycin (Sigma–Aldrich) at 37°C, 5% CO_2_, and 95% relative humidity.

### Desiccation and rehydration treatments for Pv11 cells and survivability test

Pv11 cells were subjected to desiccation-rehydration as described previously (23). Briefly, cells were incubated in a preconditioning medium (600 mM trehalose containing 10% (v/v) complete IPL-41 medium) for 48 h at 25°C. Cells were then resuspended in 400 μL of the preconditioning medium and 40 μL aliquots were dropped into 35-mm Petri dishes. The dishes were desiccated and maintained at < 10% relative humidity and 25°C for more than 10 days. After rehydration in complete IPL-41, cells were stained with propidium iodide (PI; Dojindo) and Hoechst 33342 (Dojindo) and images were acquired using a BZ-X700 microscope (Keyence). PI and Hoechst fluorescence images were acquired using a BZ-X OP-87764 microscope and an OP-87762 filter (PI: ex, 545/25 nm; em, 605/70 nm; Hoechst 33342: ex, 360/40 nm; em, 460/50 nm). The viability rate was calculated as the ratio of the number of live cells (Hoechst-positive and PI-negative) to that of total cells (Hoechst-positive).

### Vector construction for CRIS-PITCh

The CRIS-PITCh system for Pv11 cells was described in a previous study (16). The guide RNA expression vector pPvU6b-DmtRNA*-Strt1* (Supplemental Dataset S2) was constructed by replacing the gRNA region in pPvU6b-DmtRNA*-*BbsI (16) using the NEBuilder HiFi Assembly kit (New England BioLabs, Ipswich, MA, USA) as previously described (16, 17). A pair of oligonucleotides (Oligonucleotide set 1 in Supplemental Table S1) was annealed, and then ligated into the pPvU6b-DmtRNA*-* BbsI vector digested with *Bbs*I.

The donor vectors pCR4-121-AcGFP1_*Strt1* and pCR4-121-ZeoR_*Strt1* (Supplemental Dataset S2) were constructed using PCR, a HiFi Assembly kit (New England BioLabs) and a TOPO cloning kit (Thermo Fisher Scientific) as previously described (16, 17, 55). Briefly, AcGFP1 and ZeoR expression units were amplified from pPv121-AcGFP1-Pv121-ZeoR (15) as a PCR template using specific primers (Oligonucleotide set 2 in Supplemental Table S1); these amplicons contained the microhomology regions and the gRNA-target site for *Strt1*. The PCR products were cloned into the pCR4-TOPO vector (Thermo Fisher Scientific).

### Clonal *Strt1*^−/−^ cell line

A clonal knockout cell line was derived according to previous reports (16, 17). Briefly, for the establishment of the *Strt1*-knockout cell line in Figure 1C, the donor vectors harboring AcGFP1 and ZeoR expression cassettes were integrated into exon 3 of each *Strt1* allele in the opposite orientation to the target gene. The gRNA- and SpCas9-expression and donor vectors were transfected into Pv11 cells. Transfection was carried out using a NEPA21 Super Electroporator (Nepa Gene, Chiba, Japan) as described previously (16). Five days after transfection, cells were treated with 400 μg/mL Zeocin. To establish clonal cell lines, single-cell sorting was performed on a MoFlo Astrios cell sorter (Beckman Coulter, Brea, CA, USA) equipped with 355- and 488-nm lasers. One thousand wild-type Pv11 cells were seeded as a feeder layer in each well of a 96-well plate prior to cell sorting. The cells were stained with DAPI (Dojindo, Kumamoto, Japan) and DAPI and AcGFP1 were excited with 355-nm and 488-nm lasers, respectively. After single cell sorting, the cells were incubated for a while to allow them to grow, and then were treated with Zeocin to remove feeder cells. Fluorescent cell images were acquired using a BZ-X700 microscope (Keyence, Osaka, Japan). AcGFP1 fluorescence images were observed using a BZ-X GFP OP-87763 filter (AcGFP1: ex, 470/40 nm; em, 525/50 nm).

### Genomic PCR and sequencing analysis

To confirm that a precise gene knock-in was achieved in both *Strt1* alleles, genomic PCR and sequencing were performed. Genomic DNA from WT and *Strt1*^−/−^ cells was extracted with a NucleoSpin Tissue kit (Takara Bio, Shiga, Japan) and subjected to PCR using specific primers (Oligonucleotide sets 3, 4, and 5 in Supplemental Table S1). Electrophoresis images of the genomic PCR products were acquired using a ChemiDoc Touch imaging system (Bio-Rad, Hercules, CA, USA). After gel purification of the PCR products, sub-cloning was carried out using a TOPO cloning kit (Thermo Fisher Scientific), and the plasmids were subjected to Sanger sequencing. With Oligonucleotide set 3, the predicted size of the amplified products is 4,655 bp and 4,310 bp for the insertion of each of the AcGFP1 and ZeoR expression units into a *Strt1* allele, respectively, while the intact (WT) alleles should yield a 2,354 bp amplicon. With Oligonucleotide sets 4 and 5, the predicted amplicon size from the *Strt1* allele inserted with AcGFP1 and ZeoR is 583 bp and 735 bp, respectively.

### CHO-*Strt1* cell lines

The use of the Flip-In^TM^ system (Thermo Fisher Scientific) to generate a stable expression system in CHO cells was described previously (70, 71). To establish a stable CHO cell line expressing *Strt1*, we made the expression vector pcDNA5/FRT-Strt1 (Supplemental Dataset S2). The *Strt1* ORF was amplified from cDNA derived from Pv11 cells using specific primers (Oligonucleotide set 4 in Supplemental Table S1) and then cloned into the pCR4-TOPO vector. Then, to construct a Flip-In compatible expression vector for *Strt1*, the ORF in the pCR4-TOPO vector was re-amplified with specific primers (Oligonucleotide set 5 in Supplemental Table S1) and cloned into BamHI- and XhoI-digested pcDNA5/FRT vector using the NEBuilder HiFi DNA Assembly kit (New England Biolabs). The resultant construct, pcDNA5/FRT-Strt1 (Supplemental Dataset S2), was transfected with the FLP recombinase expression vector pOG44 (Thermo Fisher Scientific) into Flip-In^TM^-CHO cells. Thereafter, CHO cells constitutively expressing *Strt1* were obtained by selection with hygromycin according to the manufacturer’s manual. As a negative control, pcDNA5/FRT instead of the *Strt1* construct was transfected into Flip-In^TM^-CHO cells. The cells obtained were called CHO-X.

### Synthesis of capped RNA and *Xenopus* oocyte expression system

Use of the *Xenopus* oocyte expression system was described previously (24). An expression vector for *Strt1* was constructed from the corresponding cDNA by amplification with specific primers for *Strt1* containing restriction sites for BglII and EcoRV at the 5’-ends as the forward and reverse primers, respectively (Oligonucleotide set 6 in Supplemental Table 1). After digestion with BglII and EcoRV, the amplified cDNA was ligated into the corresponding restriction sites of the expression vector pT7XbG3 (Supplemental Dataset 2). *Strt1* cRNA was synthesized from this construct, termed pT7XbG3-*Strt1* (Supplemental Dataset 2), using the mMessage mMachine^®^ T7 kit (Thermo Fisher Scientific) according to the manufacturer’s standard protocols. The cRNAs were kept at –80°C until use. The synthesized cRNA was injected into *Xenopus* oocytes after removal of follicular cells. To express exogenous proteins, the injected oocytes were incubated in modified Barth’s saline (MBS) buffer, pH 7.8 for 3 days at 20°C. Water-injected oocytes were used to evaluate endogenous transporter activities.

### Quantification of sugars and trehalose transport activity

The methods used were described previously (24, 70, 71). To evaluate STRT1 transporter activity, *Xenopus* oocytes and CHO-cell lines were used as heterologous expression systems and were incubated in MBS and HBSS buffer containing sugars plus several other compounds, respectively. To investigate trehalose content in Pv11 cell lines such as WT and *Strt1*^−/−^ cells during preconditioning and rehydration, cells were incubated in preconditioning medium (600 mM trehalose containing 10% (v/v) complete IPL-41 medium) and complete IPL-41 medium as the rehydration medium at 25°C. To assess the NaCl-dependency of trehalose transport activity in the Pv11 cell lines, cells were incubated for 6 h in various concentrations of NaCl dissolved in complete IPL-41 medium.

The treated cells were rinsed twice with either ice-cold MBS for the oocytes or ice-cold Dulbecco’s PBS (Sigma–Aldrich) for the CHO cells, and then homogenized in 80% (v/v) EtOH containing 1% (w/v) sorbitol as an internal standard for sugar quantification. The supernatant of the homogenates was dried off with a vacuum concentrator (VC-36R, TAITEC, Saitama, Japan), and then dissolved in MilliQ water. The sugars in the samples were quantitated using an HPLC system (LC-20 system, Shimadzu, Kyoto, Japan) with a refractive index detector (RID-10A, Shimadzu). A HPX-87C column (Bio-Rad) was fitted to the HPLC with heating at 85°C to separate the sugars.

### Cell circularity

Image processing and analysis were performed using ImageJ 1.53t with Java 1.8.0_345 (64-bit) for macOS (72). Briefly, brightness and contrast were modified in bright-field images to define the cell shape. Thereafter morphology was quantified by circularity (4π × area/perimeter^2^), which corresponds to a value of 1 for a perfect circle, using ImageJ software with at least 100 cells for each condition. Detailed analysis methods, including the macros to calculate the cell circularity, are described in Supplemental procedures. All fluorescence images and original images are shown in Supplemental Fig. S4 and S5.

### Statistical analysis

All data are expressed as mean ± 95% CI (Confidence Interval) by GraphPad Prism 9 software (GraphPad, San Diego, CA, USA) was used for the statistical analyses. Differences between the two groups were examined for statistical significance using the Student *t*-test or Mann-Whitney test. Statistical significance among more than three groups was examined by ANOVA followed by Tukey’s or Dunnett’s multiple comparison test. Each statistical method is described in the Figure Legends.

### Data availability

All nucleotide sequence data in this manuscript are summarized as a GenBank-formatted file (Supplemental Dataset S2.txt). The raw data for preparation of all Figures are available as Supplemental Dataset S4.xlsx.

## Abbreviations

cRNA: capped RNA
MBS: modified Barth’s saline
CHO: Chinese hamster ovary
HBSS: Hank’s buffered saline solution
FBS: fetal bovine serum
SLC: solute carrier
SGLT: sodium/glucose co-transporter
SSS: sodium/solute symporter
TRET1: trehalose transporter 1
STRT1: sodium ion trehalose transporter 1
Treh: trehalase
G6P: glucose-6-phosphate
Hex: hexokinase
PPP: pentose-phosphate pathway
CRIS-PITCh: CRISPR/Cas9-mediated precise integration into target chromosome.

## Acknowledgments

We thank Tomoe Shiratori for maintaining Pv11 cells. We would also like to thank Clinton James Belott, whose valuable comments improved the paper. This research was partially supported by a Grant-in-Aid from Japan Society for the Promotion of Science (JSPS) KAKENHI grant numbers 23KJ0537 to KM; 12J110839 and 26850216 to SK; and 25252060 and 22H00372 to TK. The research was also supported by JST SPRING grant number JPMJSP2108 to KM.

## Notes

### Competing Interest Statement

The authors have declared no competing interest.

### Summary of Updates

Corrected author's affiliation address. Add title to Suppl. Table S1.

## References

1. J. S. Clegg, The origin of trehalose and its significance during emergence of encysted dormant embryos of *Artemia salina*. Comp Biochem Physiol B Biochem Mol Biol 14, 135–143 (1965).

2. J. S. Clegg, Cryptobiosis–a peculiar state of biological organization. Comp Biochem Physiol B Biochem Mol Biol 128, 613–624 (2001).

3. J. S. Clegg, Desiccation tolerance in encysted embryos of the animal extremophile, *Artemia*. Integr Comp Biol 45, 715–724 (2005).

4. M. Watanabe, Anhydrobiosis in invertebrates. Applied Entomology and Zoology 41, 15–31 (2006).

5. A. Adams, Cryptobiosis in Chironomidae (Diptera) – two decades on. Antenna: Bull R Entomol Soc Lond 8, 58–61 (1983).

6. H. E. Hinton, A new chironomid from Africa, the larva of which can be dehydrated without injury. Proc Zool Soc London 121, 371–380 (1951).

7. R. Cornette, T. Kikawada, The induction of anhydrobiosis in the sleeping chironomid: current status of our knowledge. IUBMB Life 63, 419–429 (2011).

8. H. E. Hinton, Cryptobiosis in the larva of *Polypedilum vanderplanki* Hint. (Chironomidae). J Insect Physiol 5, 286–288 (1960).

9. Y. Nakahara et al., Cells from an anhydrobiotic chironomid survive almost complete desiccation. Cryobiology 60, 138–146 (2010).

10. O. Gusev et al., Comparative genome sequencing reveals genomic signature of extreme desiccation tolerance in the anhydrobiotic midge. Nat Commun 5, 4784 (2014).

11. T. G. Yamada et al., Transcriptome analysis of the anhydrobiotic cell line Pv11 infers the mechanism of desiccation tolerance and recovery. Sci Rep 8, 17941 (2018).

12. T. G. Yamada et al., Identification of a master transcription factor and a regulatory mechanism for desiccation tolerance in the anhydrobiotic cell line Pv11. PLoS One 15, e0230218 (2020).

13. Y. Yoshida, et al., High quality genome assembly of the anhydrobiotic midge provides insights on a single chromosome-based emergence of extreme desiccation tolerance. NAR Genom Bioinform 4, lqac029 (2022).

14. Y. Sogame et al., Establishment of gene transfer and gene silencing methods in a desiccation-tolerant cell line, Pv11. Extremophiles 21, 65–72 (2017).

15. Y. Miyata et al., Identification of a novel strong promoter from the anhydrobiotic midge, *Polypedilum vanderplanki*, with conserved function in various insect cell lines. Sci Rep 9, 7004 (2019).

16. Y. Miyata et al., Cas9-mediated genome editing reveals a significant contribution of calcium signaling pathways to anhydrobiosis in Pv11 cells. Sci Rep 11, 19698 (2021).

17. S. Tokumoto et al., Genome-wide role of HSF1 in transcriptional regulation of desiccation tolerance in the anhydrobiotic cell line, Pv11. Int J Mol Sci 22 (2021).

18. M. Watanabe, T. Kikawada, N. Minagawa, F. Yukuhiro, T. Okuda, Mechanism allowing an insect to survive complete dehydration and extreme temperatures. J Exp Biol 205, 2799–2802 (2002).

19. K. A. C. Madin, J. H. Crowe, Anhydrobiosis in nematodes: Carbohydrate and lipid metabolism during dehydration. J Exp Zool 193, 335–342 (1975).

20. G. M. Gadd, K. Chalmers, R. H. Reed, The role of trehalose in dehydration resistance of Saccharomyces cerevisiae. FEMS Microbiol Lett 48, 249–254 (1987).

21. J. H. Crowe, L. M. Crowe, D. Chapman, Preservation of membranes in anhydrobiotic organisms: the role of trehalose. Science 223, 701–703 (1984).

22. M. Sakurai et al., Vitrification is essential for anhydrobiosis in an African chironomid, *Polypedilum vanderplanki*. Proc Natl Acad Sci U S A 105, 5093–5098 (2008).

23. K. Watanabe, S. Imanishi, G. Akiduki, R. Cornette, T. Okuda, Air-dried cells from the anhydrobiotic insect, *Polypedilum vanderplanki*, can survive long term preservation at room temperature and retain proliferation potential after rehydration. Cryobiology 73, 93–98 (2016).

24. T. Kikawada et al., Trehalose transporter 1, a facilitated and high-capacity trehalose transporter, allows exogenous trehalose uptake into cells. Proc Natl Acad Sci U S A 104, 11585–11590 (2007).

25. Y. Kanamori et al., The trehalose transporter 1 gene sequence is conserved in insects and encodes proteins with different kinetic properties involved in trehalose import into peripheral tissues. Insect Biochem Mol Biol 40, 30–37 (2010).

26. M. Watanabe, T. Kikawada, T. Okuda, Increase of internal ion concentration triggers trehalose synthesis associated with cryptobiosis in larvae of *Polypedilum vanderplanki*. J Exp Biol 206, 2281–2286 (2003).

27. E. M. Wright, D. D. Loo, B. A. Hirayama, Biology of human sodium glucose transporters. Physiol Rev 91, 733–794 (2011).

28. G. Gyimesi, J. Pujol-Gimenez, Y. Kanai, M. A. Hediger, Sodium-coupled glucose transport, the SLC5 family, and therapeutically relevant inhibitors: from molecular discovery to clinical application. Pflugers Arch 472, 1177–1206 (2020).

29. M. M. Ceder, R. Fredriksson, A phylogenetic analysis between humans and *D. melanogaster*: A repertoire of solute carriers in humans and flies. Gene 809, 146033 (2022).

30. I. S. Wood, P. Trayhurn, Glucose transporters (GLUT and SGLT): expanded families of sugar transport proteins. Br J Nutr 89, 3–9 (2003).

31. B. Thorens, M. Mueckler, Glucose transporters in the 21st Century. Am J Physiol Endocrinol Metab 298, E141–145 (2010).

32. E. M. Wright, Glucose transport families SLC5 and SLC50. Mol Aspects Med 34, 183–196 (2013).

33. S. M. Denecke et al., The Identification and Evolutionary Trends of the Solute Carrier Superfamily in Arthropods. Genome Biol Evol 12, 1429–1439 (2020).

34. S. P. Alexander et al., THE CONCISE GUIDE TO PHARMACOLOGY 2021/22: Transporters. Br J Pharmacol 178 Suppl 1, S412–S513 (2021).

35. O. Vitavska, H. Wieczorek, The SLC45 gene family of putative sugar transporters. Mol Aspects Med 34, 655–660 (2013).

36. L. Q. Chen et al., Sugar transporters for intercellular exchange and nutrition of pathogens. Nature 468, 527–532 (2010).

37. M. Mueckler, B. Thorens, The SLC2 (GLUT) family of membrane transporters. Mol Aspects Med 34, 121–138 (2013).

38. A. Diez-Sampedro et al., A glucose sensor hiding in a family of transporters. Proc Natl Acad Sci U S A 100, 11753–11758 (2003).

39. L. Bianchi, A. Diez-Sampedro, A single amino acid change converts the sugar sensor SGLT3 into a sugar transporter. PLoS One 5, e10241 (2010).

40. D. D. Loo et al., Passive water and ion transport by cotransporters. J Physiol 518, 195–202 (1999).

41. E. M. Wright, D. D. Loo, B. A. Hirayama, E. Turk, Surprising versatility of Na^+^-glucose cotransporters: SLC5. Physiology (Bethesda) 19, 370–376 (2004).

42. Y. Zhang, Y. Zhang, K. Sun, Z. Meng, L. Chen, The SLC transporter in nutrient and metabolic sensing, regulation, and drug development. J Mol Cell Biol 11, 1–13 (2019).

43. W. Song, D. Li, L. Tao, Q. Luo, L. Chen, Solute carrier transporters: the metabolic gatekeepers of immune cells. Acta Pharm Sin B 10, 61–78 (2020).

44. S. Kikuta, Y. Hagiwara-Komoda, H. Noda, T. Kikawada, A novel member of the trehalose transporter family functions as an H^+^-dependent trehalose transporter in the reabsorption of trehalose in Malpighian tubules. Front Physiol 3, 290 (2012).

45. M. Iwata, M. Yoshinaga, K. Mizutani, T. Kikawada, S. Kikuta, Proton gradient mediates hemolymph trehalose influx into aphid bacteriocytes. Arch Insect Biochem Physiol 112, e21971 (2023).

46. E. McMullen, A. Weiler, H. M. Becker, S. Schirmeier, Plasticity of carbohydrate transport at the blood-brain barrier. Front Behav Neurosci 14, 612430 (2020).

47. Y. Y. Yuan et al., Functional characterization of a novel, highly expressed ion-driven sugar antiporter in the thoracic muscles of *Helicoverpa armigera*. Insect Sci 29, 78–90 (2022).

48. D. E. Featherstone, Glial solute carrier transporters in *Drosophila* and mice. Glia 59, 1351–1363 (2011).

49. K. Stergiopoulos, P. Cabrero, S. A. Davies, J. A. Dow, Salty dog, an SLC5 symporter, modulates *Drosophila* response to salt stress. Physiol Genomics 37, 1–11 (2009).

50. M. Dus, M. Ai, G. S. Suh, Taste-independent nutrient selection is mediated by a brain-specific Na^+^ /solute co-transporter in *Drosophila*. Nat Neurosci 16, 526–528 (2013).

51. Y. Li, W. Wang, H. Y. Lim, *Drosophila* Solute Carrier 5A5 Regulates Systemic Glucose Homeostasis by Mediating Glucose Absorption in the Midgut. Int J Mol Sci 22 (2021).

52. K. Yildirim et al., Redundant functions of the SLC5A transporters Rumpel, Bumpel, and Kumpel in ensheathing glial cells. Biol Open 11 (2022).

53. T. Sakuma, S. Nakade, Y. Sakane, K. T. Suzuki, T. Yamamoto, MMEJ-assisted gene knock-in using TALENs and CRISPR-Cas9 with the PITCh systems. Nat Protoc 11, 118–133 (2016).

54. M. Raja, R. K. Kinne, Identification of phlorizin binding domains in sodium-glucose cotransporter family: SGLT1 as a unique model system. Biochimie 115, 187–193 (2015).

55. Y. Miyata et al., Identification of Genomic Safe Harbors in the Anhydrobiotic Cell Line, Pv11. Genes (Basel) 13 (2022).

56. L. Michaelis, M. L. Menten, K. A. Johnson, R. S. Goody, The original Michaelis constant: translation of the 1913 Michaelis-Menten paper. Biochemistry 50, 8264–8269 (2011).

57. E. Klipp, B. Nordlander, R. Kruger, P. Gennemark, S. Hohmann, Integrative model of the response of yeast to osmotic shock. Nat Biotechnol 23, 975–982 (2005).

58. C. Saldana et al., Rapid and reversible cell volume changes in response to osmotic stress in yeast. Braz J Microbiol 52, 895–903 (2021).

59. F. Lang et al., Functional significance of cell volume regulatory mechanisms. Physiol Rev 78, 247–306 (1998).

60. P. H. Yancey, Water Stress, Osmolytes and Proteins. Am Zool 41, 699–709 (2001).

61. R. D. Sleator, C. Hill, Bacterial osmoadaptation: the role of osmolytes in bacterial stress and virulence. FEMS Microbiol Rev 26, 49–71 (2002).

62. C. Ziegler, E. Bremer, R. Kramer, The BCCT family of carriers: from physiology to crystal structure. Mol Microbiol 78, 13–34 (2010).

63. S. M. Hosseiniyan Khatibi et al., Osmolytes resist against harsh osmolarity: Something old something new. Biochimie 158, 156–164 (2019).

64. A. Ryabova et al., Combined metabolome and transcriptome analysis reveals key components of complete desiccation tolerance in an anhydrobiotic insect. Proc Natl Acad Sci U S A 117, 19209–19220 (2020).

65. K. Mitsumasu et al., Enzymatic control of anhydrobiosis-related accumulation of trehalose in the sleeping chironomid, *Polypedilum vanderplanki*. FEBS J 277, 4215–4228 (2010).

66. B. Li, C. N. Dewey, RSEM: accurate transcript quantification from RNA-Seq data with or without a reference genome. BMC Bioinform 12, 323 (2011).

67. M. G. Grabherr et al., Full-length transcriptome assembly from RNA-Seq data without a reference genome. Nat Biotechnol 29, 644–652 (2011).

68. S. R. Eddy, Accelerated Profile HMM Searches. PLoS Comput Biol 7, e1002195 (2011).

69. T. Paysan-Lafosse et al., InterPro in 2022. Nucleic Acids Res 51, D418–D427 (2023).

70. T. Uchida, M. Furukawa, T. Kikawada, K. Yamazaki, K. Gohara, Intracellular trehalose via transporter TRET1 as a method to cryoprotect CHO-K1 cells. Cryobiology 77, 50–57 (2017).

71. T. Uchida, M. Furukawa, T. Kikawada, K. Yamazaki, K. Gohara, Viabilities of long-term cryopreserved CHO-TRET1 cells with trehalose and DMSO. Bull Glaciol Res 37, 1–9 (2019).

72. C. A. Schneider, W. S. Rasband, K. W. Eliceiri, NIH Image to ImageJ: 25 years of image analysis. Nat Methods 9, 671–675 (2012).

