## Supplemental information for "A Sodium-dependent Trehalose Transporter Contributes to Anhydrobiosis in Insect Cell Line, Pv11"

**This PDF file includes:**

Supporting information text
SI References
Supplemental procedures
Figures S1 to S6

**Other supporting materials for this manuscript include the following:**

Table S1
Datasets S1 to S4

**Supporting Information Text**

To address the physiological role of TRET1, a high capacity and facilitated diffusion-type trehalose transporter (1) in Pv11 cells, we attempted to knock out *Tret1* using the same method used to generate *Strt1*$^{-/-}$. However, so far no *Tret1*$^{-/-}$ knockout cells have been established. In our method, knockout cell lines are generated by inserting expression units of genes encoding selection markers, such as antibiotic resistance or fluorescent proteins, in the opposite orientation to that of the target, thereby causing loss of function of the target gene. In this case, the gene expression units encoding zeocin resistance and AcGFP1 were respectively inserted into each allele for *Tret1*, so we expected to select for AcGFP1-fluorescent cells that proliferate in Zeocin-containing medium. However, when attempting to generate *Tret1*-knockout cells, we were not able to obtain cells that survived after zeocin treatment, even after multiple repetitions. Pv11-KH cells, which have already been established, permanently express AcGFP1 but do not die in normal culture (2). Hence, the death of *Tret1*$^{-/-}$ cells after Zeocin treatment must not be due to cytotoxicity caused by the constitutive expression of AcGFP1. Therefore, we reasoned that *Tret1* knockout might have a lethal effect on Pv11 cells because TRET1 might contribute to the normal proliferation of cells.

The Migdebase2 database shows that *Tret1* (gene ID: g6534) is highly expressed under normal culture conditions prior to desiccation in Pv11 cells (3). Thus, TRET1 might be involved in the proliferation and maintenance of survival of Pv11 cells not only during anhydrobiosis but also during normal growth. The IPL-41 culture medium used to culture Pv11 cells contains sugars such as (with approximate concentrations) 13.8 mM D-glucose, 4.8 mM sucrose and 2.9 mM maltose as nutrient sources. TRET1 can effectively transport glucose as well as trehalose, and also sucrose and maltose to some extent (1). Therefore, in Pv11 cells, TRET1 functions to take up sugars as nutrients from the medium.

These results suggest that TRET1 is involved in cell proliferation and survival by taking up glucose and sucrose from the medium for use as a carbon and energy source during normal growth. Thus, inactivating *Tret1* would cause cell lethality. In fact, in *Drosophila*, complete deletion of two paralogs of *Tret1* results in early larval lethality (4), suggesting that TRET1 is an essential transporter for normal growth in insects. To overcome the issue of the lethality of *Tret1* knockout, we will apply RNAi technology to establish a method of suppressing TRET1 expression during the desiccation process only, without changing the transcription level of *Tret1* during desiccation.

**Supplemental procedures**

**Outline of analysis for cell circularity**

1. Start ImageJ

2. Check "Shape descriptors" in "Set measurement.

3. Run the macro: In ImageJ go to "*Plugins*" and then "*New*", "*Macro*".

4. <A-1> pipeline was used to evaluate the cells during trehalose preconditioning (Fig. 5A and Supplemental Fig. S4). Alternatively, <A-2> pipeline was used to do the cells after rehydration (Fig. 5B and Supplemental Fig. S5).

5. Open the fluorescent image file and press Run.

**Macros**

The basic parameters for the code development were developed using the macro record function. "Plugins>Macros>Record". A basic outline of the code was used, following existing ImageJ protocols. The code is designed to work on an image stack of AcGFP1 fluorescent cell images of single channel.

In this macro, the cell contour extraction is conducted using a succession of classical ImageJ functions. The stack was first converted to 8-bit. The signal was then binarized using the "Threshold" function, and separate background and cells. Cells above a certain size were recognized and the shape of the cells was measured. The thresholds were set to consider differences in extracellular osmotic conditions during trehalose preconditioning and rehydration.

<A-1> A macro pipeline for trehalose preconditioning process. This pipeline was applied for images in Supplemental Fig. S4.

```
run("8-bit");

setAutoThreshold("Default dark no-reset");

//run("Threshold...");
```

```
setThreshold(3, 255, "raw");

run("Analyze Particles...", "size=100-Infinity pixel show=Outlines display clear overlay add");
```

<A-2> A macro pipeline for rehydration process. This pipeline was applied for images in Supplemental Fig. S5.

```
run("8-bit");

setAutoThreshold("Default dark no-reset");

//run("Threshold...");

setThreshold(20, 255, "raw");

run("Analyze Particles...", "size=100-Infinity pixel show=Outlines display clear overlay add");
```

**SI References**


1. T. Kikawada *et al.*, Trehalose transporter 1, a facilitated and high-capacity trehalose transporter, allows exogenous trehalose uptake into cells. *Proc Natl Acad Sci U S A* **104**, 11585-11590 (2007).

2. Y. Sogame *et al.*, Establishment of gene transfer and gene silencing methods in a desiccation-tolerant cell line, Pv11. *Extremophiles* **21**, 65-72 (2017).

3. Y. Yoshida *et al.*, High quality genome assembly of the anhydrobiotic midge provides insights on a single chromosome-based emergence of extreme desiccation tolerance. *NAR Genom Bioinform* **4**, lqac029 (2022).

4. A. Volkenhoff *et al.*, Glial Glycolysis Is Essential for Neuronal Survival in *Drosophila*. *Cell Metab* **22**, 437-447 (2015).

### Supplemental Figures, Table and Datasets

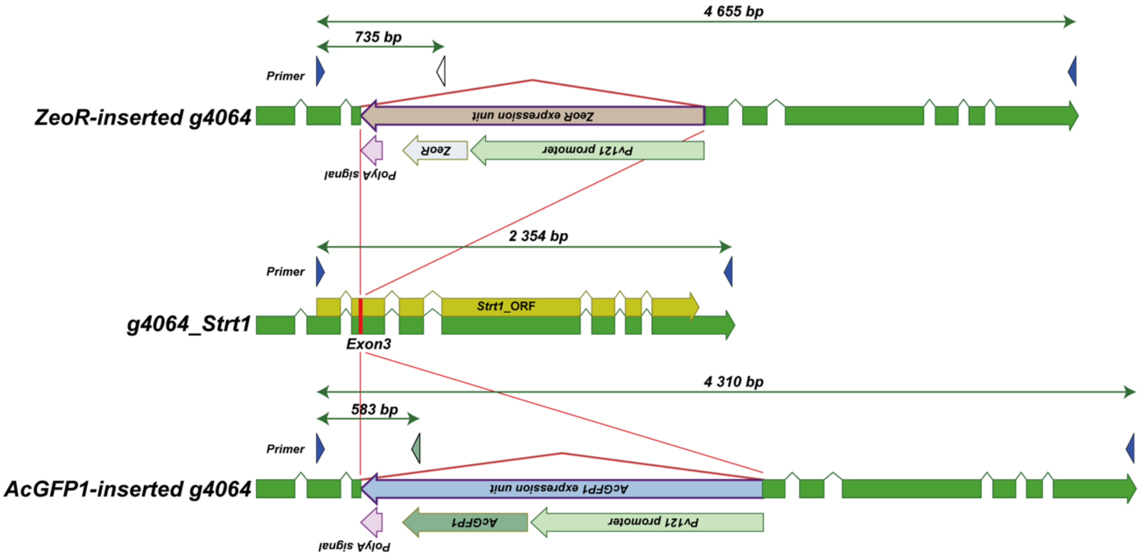


**Fig. S1. Detailed schema of structures of intact and knocked-in *Strt1* alleles**

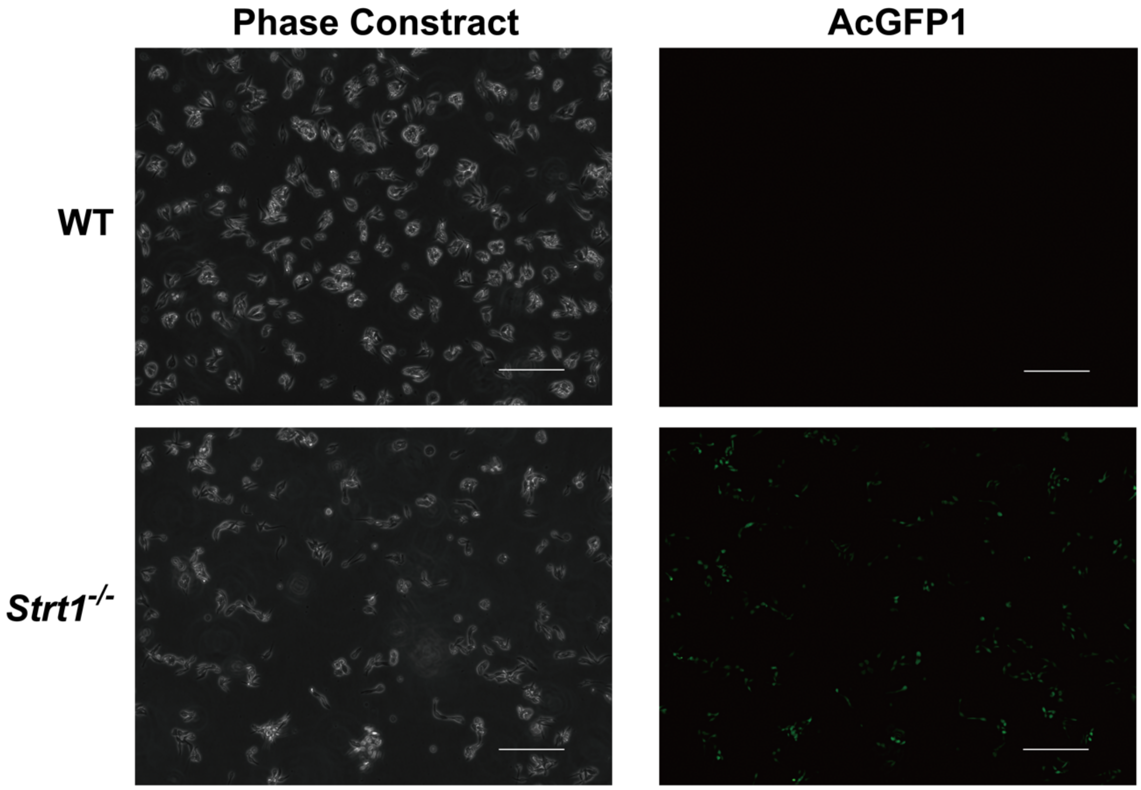


**Fig. S2. Phase contrast of Pv11 WT and *Strt1*$^{-/-}$ cells.**

121-AcFFP1 sequence was knocked into exon3 of *Strt1*, as confirmed by fluorescence images. Scale bar represents 100 µm. *Strt1*$^{-/-}$ cells show AcGFP1 fluorescence, but wild-type Pv11 cells (WT) do not.

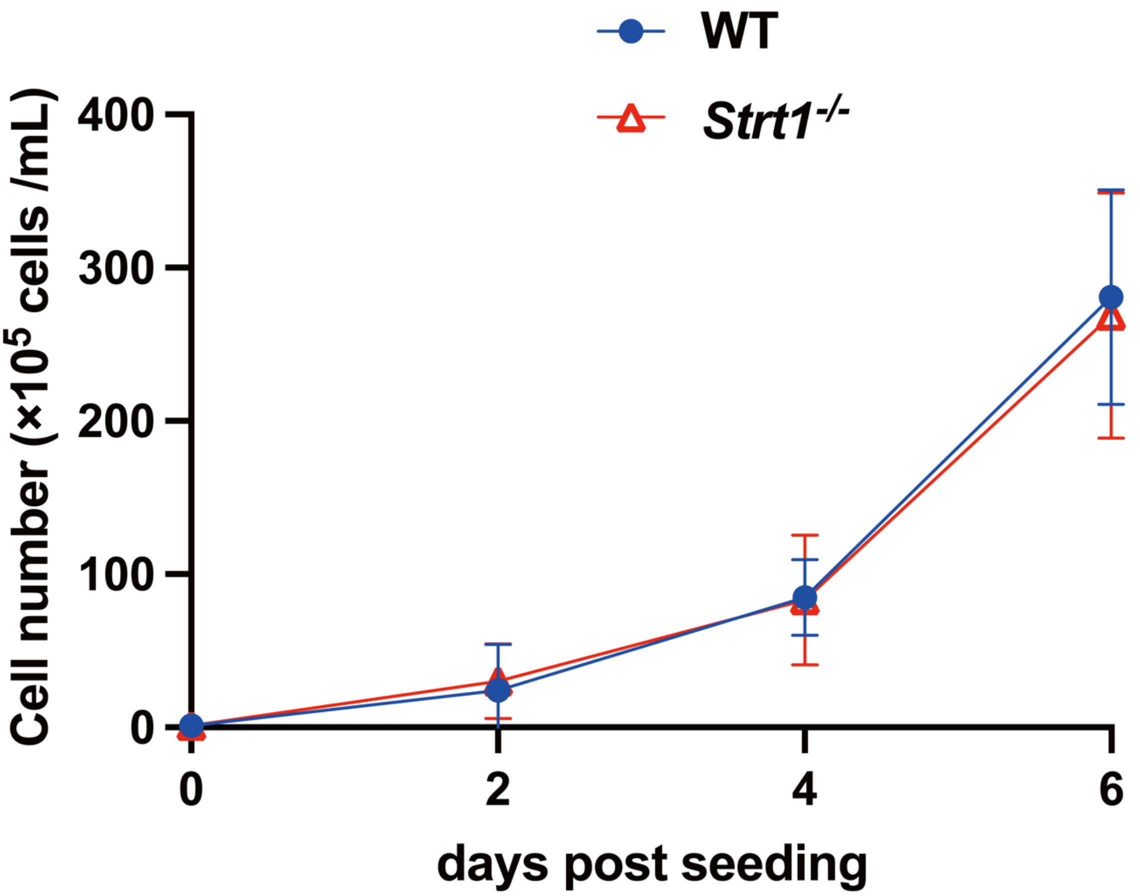


**Fig. S3. Cell proliferation rate in culture without Zeocin.**

The proliferation rates of the WT and *Strt1*$^{-/-}$ cell lines are shown. The proliferation rates of both cell lines are essentially identical. Values are expressed as mean ± 95 % CI; n =3 in each group.

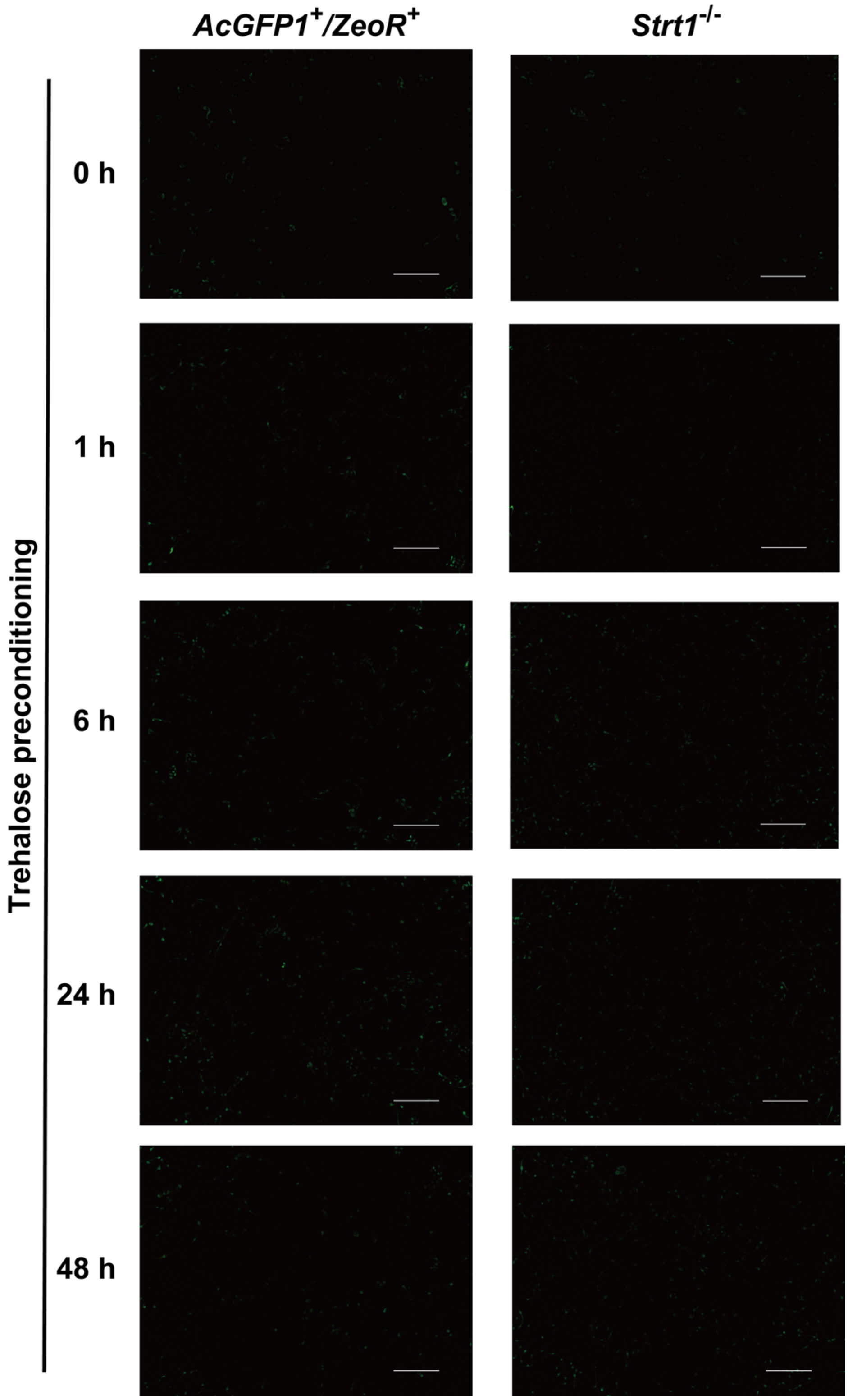


**Fig. S4. Cell morphology during trehalose preconditioning.**

Images of cells during trehalose pre-conditioning were obtained to determine cell circularity, which was then quantified using ImageJ. Scale bar represents 100 μm.

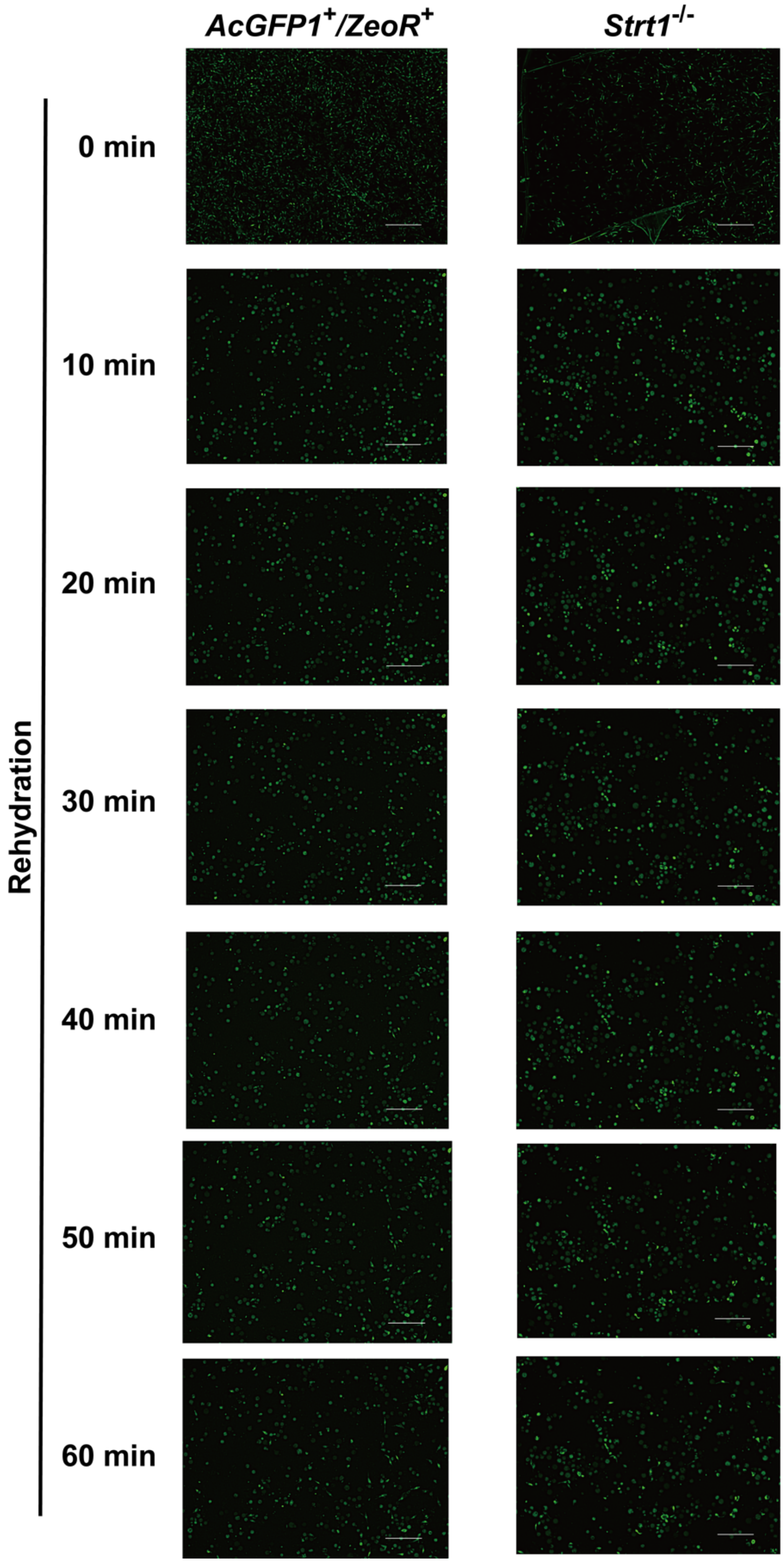


**Fig. S5. Cell morphology after rehydration.**

Images of cells during rehydration were obtained to determine cell circularity, which was quantified using ImageJ. Scale bar represents 100 μm.

A

B

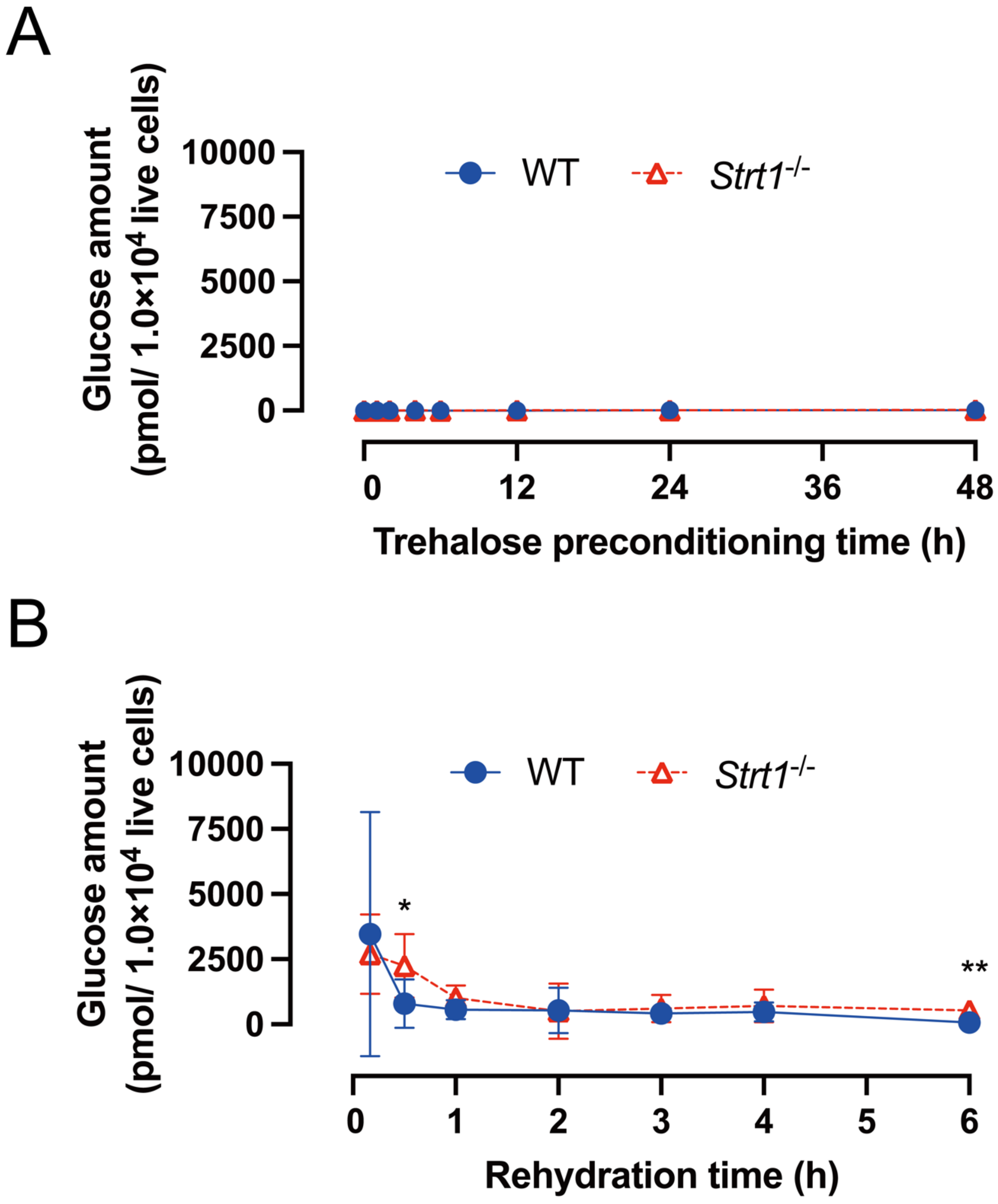


**Fig. S6. Comparison of glucose content in *Strt1*$^{-/-}$ and WT cells.**

(A) Changes in glucose content in cells during preconditioning with 600 mM trehalose solution as the induction step for anhydrobiosis. (B) Changes in glucose content in dried cells after rehydration. *: $p \leq 0.05$, **: $p \leq 0.01$. Values are expressed as mean ± 95 % CI; n =4 in each group.

**Table S1. Primers for construction of the donor vectors used for construction of gRNA-expression vectors and for genomic DNA sequencing.**

**Dataset S1. Expression profiles of all SLC5 family genes during preconditioning and rehydration of Pv11 cells.**

Transcriptome data for 22 genes encoding proteins with SLC5 (SSF)-specific motifs were retrieved from the latest genome database for Pv11 cells, Midgebase2 (https://www.midgebase.org). Each value indicates TPM.

**Dataset S2. Nucleotide sequences.**

Sequence information for the plasmids, *Strt1* genome and cDNA, and the alleles for *Strt1* knockout by CRIS-PITCh, described in this paper, was compiled as GenBank-formatted text.

**Dataset S3. Expression of genes for carbohydrate metabolism during anhydrobiosis of Pv11 cells.**

Transcriptome data for TRET1 (*g6534)*, trehalase (*g17178*) and hexokinases (*g2925* and *g1897*) were obtained from Midgebase2 (https://www.midgebase.org). Each value indicates TPM.

**Dataset S4. Raw data for experiments in Figures.**

The dataset contains the experimental values and statistical results used in the figures.
